# AUF1-Engineered Intestinal Organoids Enhance Epithelial Barrier Repair and Mucosal Regeneration in Experimental Colitis

**DOI:** 10.64898/2026.08.21.746163

**Authors:** Oishika Das, Sutanu Acharya Chowdhury, Animesh Gope, Anita Nanda Goswami, Moumita Bhaumik

## Abstract

Inflammatory bowel disease (IBD) often involves disrupted intestinal epithelial barrier, but therapies specifically targeting this barrier are limited. We found that downregulated AUF1 (HNRNPD) contributes to defective barrier integrity in ulcerative colitis (UC). Compared to controls, its expression level was decreased and inversely correlated with clinical severity. Knocking down AUF1 in human and mouse colonic organoids led to impaired barrier function, with reduced Occludin and upregulated Claudin-2, mimicking characteristic IBD-associated mucosal alterations. Distinct RNA-binding activity of AUF1 protein isoforms contributed to these changes: p37 stabilized Occludin mRNA and blocked microRNA-122/Ago2-mediated repression, whereas p40 promoted Claudin-2 mRNA degradation via ubiquitin-proteasome pathway. Restoring AUF1 expression in organoids enhanced epithelial properties and, when transplanted into mice with established colitis, accelerated mucosal healing and epithelial regeneration in recipient mice and decreased fibrosis. Our study unravelled a post-transcriptional mechanism important for intestinal homeostasis and demonstrated a concept of using engineered organoids for treating IBD.

## Introduction

Inflammatory bowel disease (IBD), encompassing Crohn’s disease and ulcerative colitis, is characterized by chronic relapsing inflammation that progressively erodes epithelial barrier integrity and compromises mucosal homeostasis [1]. Although many therapies are currently targeting immune processes, a significant number of patients cannot achieve long-lasting remission, and there is a great need to develop approaches that directly restore epithelial integrity. Indeed, mucosal healing, which refers to the complete recovery of the epithelium, is recognized as a key player in long-term outcome in IBD [2].

Organoids represent a promising regenerative medicine approach for the treatment of IBD. This therapy is based on the fact that intestinal organoids can be isolated from stem cells, expanded in vitro, and then used for transplantation. Although there is preliminary evidence that the transplantation of organoids can induce epithelial repair, organoids often fail to demonstrate long-lasting efficacy, and it is unclear how their function can be optimized to achieve durable restoration of the epithelium.

The fact is that the intestinal epithelium itself regulates mucosal immunity and integrity, which means promoting its regeneration could have a significant impact on the outcome of IBD [3]. Tight junctions (TJ) are protein complexes located at the apical surface of intestinal epithelial cells that regulate the paracellular permeability barrier [4]. Epithelial cell death, TJ disruption, and subsequent loss of barrier integrity are recognized as critical events in the pathogenesis of IBD [5]. Specifically, pro-inflammatory mediators, which are increased in IBD, can cause TJ instability, epithelial cell death, and even ulceration [6].

A variety of transmembrane proteins constitute tight junctions (TJs) together with occludin, tricellulin, different claudins, and junctional adhesion molecules (JAMs) [7]. As an essential component of the tight junction strand, occludin plays a significant role in regulating the permeability of epithelia and endothelia [8]. Being a part of TJs, the protein helps maintain barrier functions, contributing to the regulation of paracellular permeability. What is more, Claudin-2 is reported to be responsible for the formation of cation-selective and large-conductance aquaporin water channels in “leaky” epithelia, which are characteristic of the kidney, intestine, and other body parts [9]. The decrease in occludin with the concomitant increase in Claudin-2 expression was found to be associated with the onset and progression of IBD [10]; [11].

A substantial cohort of TJ proteins are encoded by mRNAs harboring AU-rich elements (AREs) within their 3′ untranslated regions (3’ UTR). These motifs, via interplay with ARE-binding proteins, determines both the stability and translational fate of their transcripts [12]. One such effector is heterogeneous nuclear ribonucleoprotein D (HNRNPD), more commonly known as AU-rich element/poly(U)-binding degradation factor 1 (AUF1). AUF1 is important in modulating ARE-mRNA stability [12]. Through alternative splicing, the AUF1 transcript yields four isoforms—p37, p40, p42, and p45—each endowed with RNA-binding property, albeit with divergent affinities [12]. Historically, AUF1 has been canonized as a destabilizer of mRNAs through the assembly of ribonucleoprotein complexes [12]. Intriguingly, emerging investigations delineate a stabilizing capacity of AUF1 toward select transcripts [13], underscoring its dualistic and enigmatic comportment, which remains largely unrevealed.

Our prior inquiries revealed that ablation of AUF1 causes colitis in murine models, accentuating its physiological gravitas. Nonetheless, the precise molecular underpinnings by which AUF1 dictates epithelial barrier function remain elusive. This lacuna impels the pressing interrogation of whether AUF1 downregulation directly engenders barrier dysfunction within the intestinal epithelium.

In this study, we uncover a crucial regenerative function of AUF1 in safeguarding epithelial homeostasis and mucosal integrity. Silencing AUF1 using cell-penetrating morpholinos disrupted epithelial barrier function in both murine and human colonic organoids, closely mirroring the pathological features of colitis. Mechanistically, AUF1 binds to the mRNAs of the tight junction proteins Occludin and Claudin-2, stabilizing the former while facilitating the degradation of the latter, thereby fine-tuning epithelial permeability. Remarkably, transplantation of AUF1-expressing organoids into colitic mice not only restored epithelial structure and barrier function but also conferred long-lasting protection against mucosal inflammation. In contrast, transplantation of unmodified organoids offered only transient relief. These findings underscore the therapeutic superiority of AUF1-reconstituted organoid transplantation as a regenerative intervention capable of durable epithelial restoration and sustained remission in inflammatory bowel disease.

## Results

### AUF1 suppression correlates with compromised intestinal epithelial barrier integrity in inflammatory bowel disease

To identify key drivers of epithelial barrier dysfunction in inflammatory bowel disease, we first examined the expression of tight junction (TJ) genes in ulcerative colitis (UC) patients and healthy controls. Transcriptomic analysis revealed the most pronounced changes in two TJ components, *OCLN* and *CLDN2*: compared with healthy controls (n = 4), UC patients (n = 5) exhibited reduced *OCLN* and increased *CLDN2* expression (**Fig. 1A**). These findings were independently validated by quantitative qRT-PCR in a larger cohort (10 UC patients and 8 healthy controls; **Fig. 1B, C**). Consistent with the transcriptional changes, immunohistochemical (IHC) analysis showed reduced Occludin and increased Claudin-2 protein levels in the epithelial layer of colonic biopsies of control and IBD patients (**Fig. 1D, E**).

**Figure 1.**
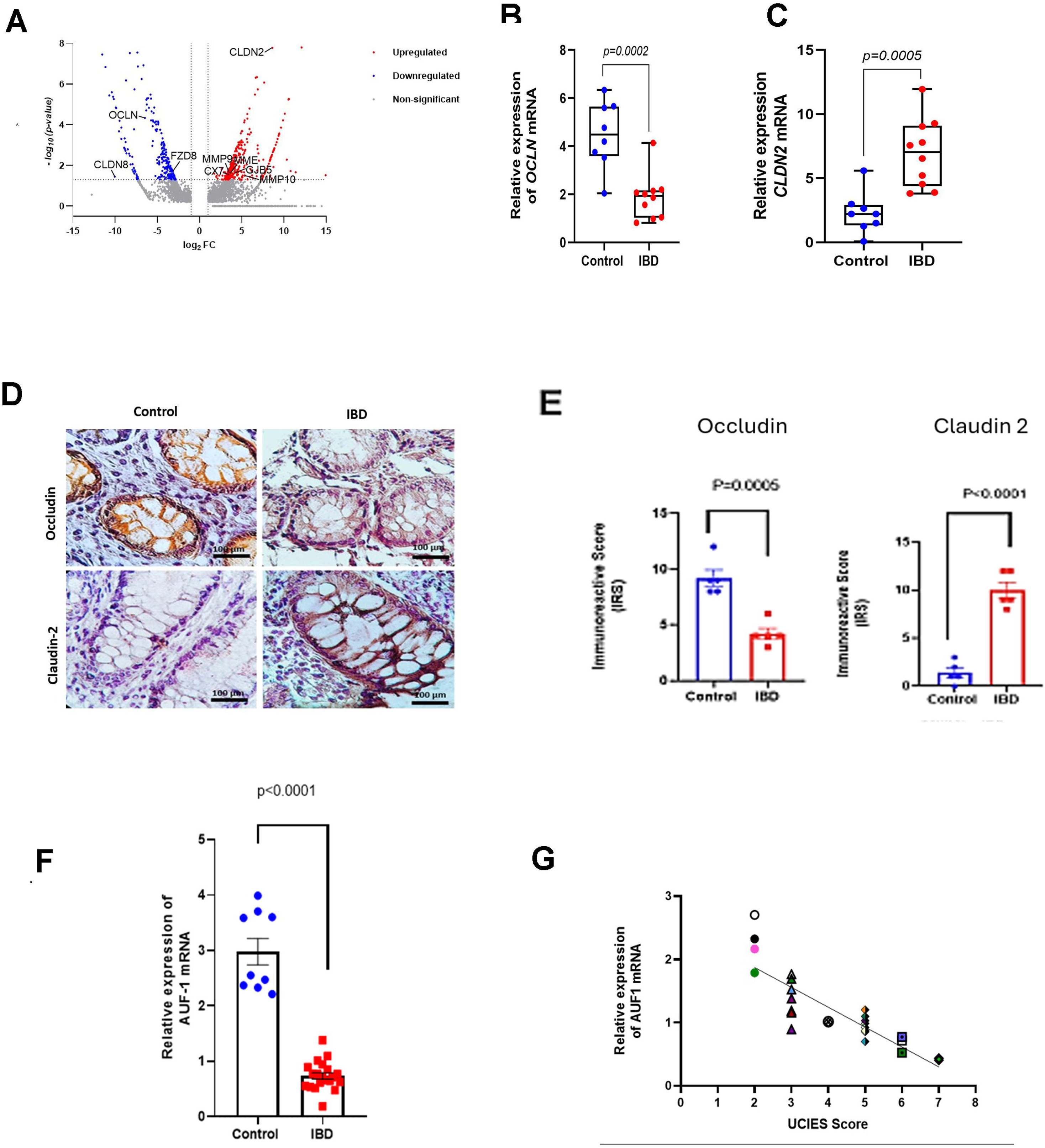
Altered tight-junction protein expression and AUF1 in inflammatory bowel disease. (A) Volcano plot showing differentially expressed genes between colonic biopsies of control and IBD patients. Upregulated genes are shown in red, downregulated genes in blue, and non-significant genes in gray; selected tight-junction–related genes are annotated. (B–C) Relative mRNA expression of Occludin (**OCLN)** (B) and Claudin 2 (**CLDN2)** (C) in control and IBD patient biopsies. Each dot represents an individual sample; box plots show the median and interquartile range. (D) Representative immunohistochemical staining for occludin and claudin-2 in control and IBD patient tissues. Scale bars, 100 μm. (E) Quantification of occludin and claudin-2 immunoreactive scores (IRS) in control and IBD patient tissues. (F) Relative mRNA expression of **AUF1** in control and IBD patient biopsies. (G) Correlation between AUF1 expression (ΔCt value) and UCIES score. Spearman’s rank correlation analysis was performed (ρ = −0.8892). Statistical significance is indicated in the individual panels.

Most TJ transcripts, including *Occludin* and *claudin 2*, contain AU-rich elements (AREs) within their 3′ untranslated regions, which serve as binding sites for RNA-binding proteins that regulate transcript stability and turnover. We therefore examined the expression of ARE-binding proteins and found that the most significantly downregulated gene in UC was *HNRNPD*, which encodes the RNA-binding protein AUF1. Reduced *AUF1* expression was validated at the mRNA level in an independent cohort of 21 UC patients and 9 healthy controls (p < 0.0001; **Fig. 1F**). Importantly, AUF1 expression exhibited a strong inverse correlation with disease severity (Spearman’s ρ = −0.8892, P = 0.0182), indicating that progressive loss of AUF1 is closely associated with epithelial barrier disruption during intestinal inflammation (**Fig. 1G**). Together, these findings associate with reduced and enhanced expression of the TJ protein, Occludin and claudin 2 respectively and downregulation of the post-transcriptional regulator AUF1, suggesting a potential link between altered RNA stability and TJ disruption in UC.

### AUF1 Loss Promotes Tight Junction Dysregulation and Intestinal Epithelial Barrier Dysfunction in patient derived organoid

To determine whether AUF1 contributes to increased epithelial permeability in the colon, we established colonic organoids from colorectal biopsies obtained from controls and IBD patients. Biopsy-derived tissues were propagated for 10 days to allow organoid formation. Organoids derived from controls were subsequently treated for 48 h with either a cell-penetrating morpholino targeting AUF1 (AUF1-MO) or a scrambled morpholino (S-MO). Treatment with 25 nM (0.025 µM) AUF1-MO did not affect organoid viability (**Fig. S1A, B**) but resulted in efficient AUF1 knockdown (**Fig. S1C**).

We next assessed AUF1 protein expression by immunofluorescence in S-MO-treated control organoids (S-MO organoid), AUF1-MO-treated organoids (AUF1-MO organoid), and organoids derived from IBD biopsies (IBD-organoid). AUF1 expression was markedly reduced in IBD-organoids compared with S-MO-organoids. As expected, AUF1-MO organoid showed reduced AUF1 expression relative to S-MO organoid indicating optimal knockdown (**Fig. 2A, B**). To assess epithelial barrier integrity, organoids were exposed to FITC-dextran (4 kDa; FITC-D4) 72 h after morpholino treatment. In S-MO-organoids, FITC-D4 fluorescence was predominantly restricted to the extraluminal compartment, resulting in a low intraluminal-to-extraluminal (IN/OUT) fluorescence ratio. In contrast, both AUF1-MO-organoids and IBD-organoids showed increased accumulation of FITC-D4 within the lumen, reflected by a significantly higher IN/OUT fluorescence ratio (**Fig. 2C, D**).

**Figure 2.**
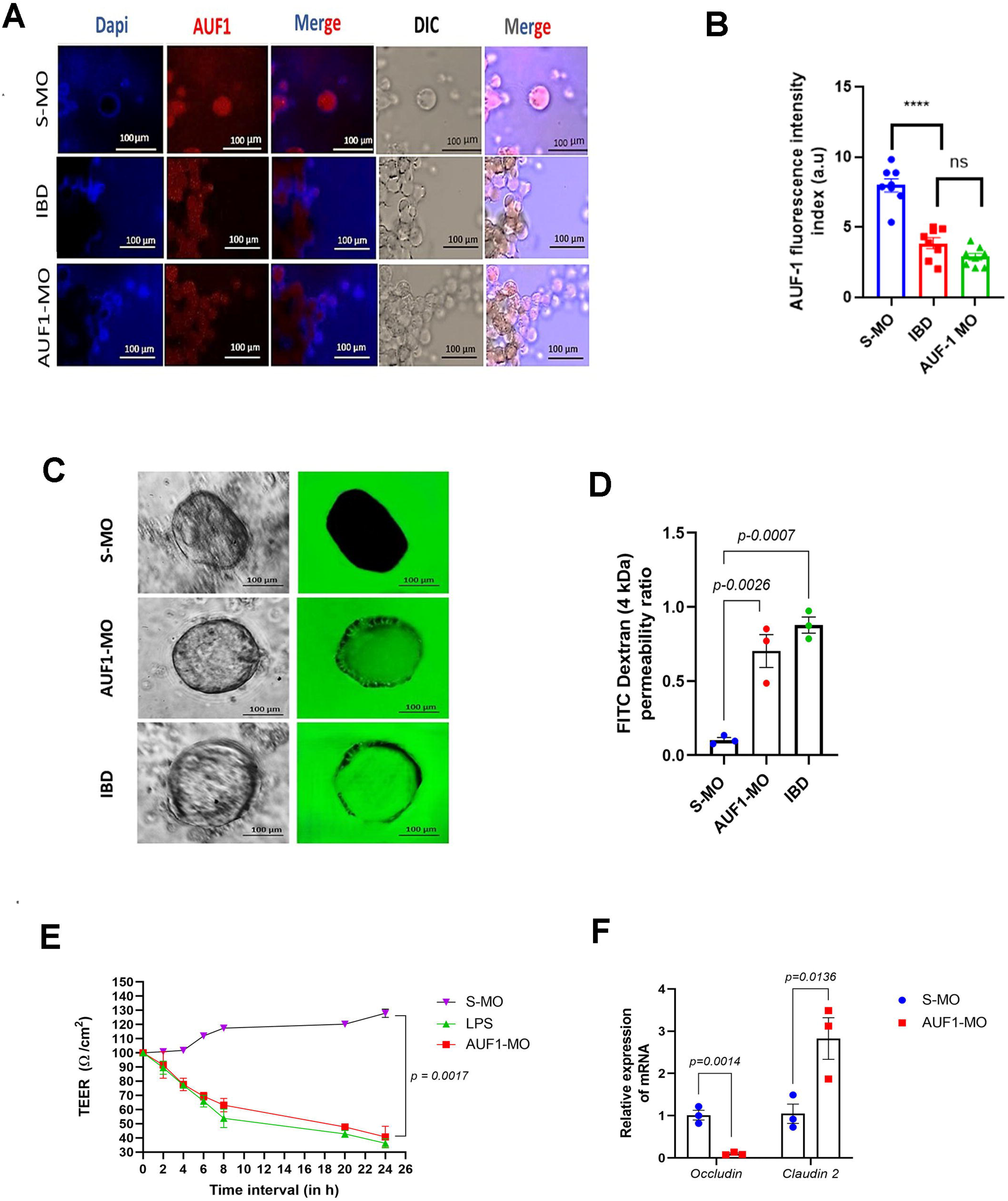
AUF1 deficiency impairs intestinal epithelial barrier integrity patient derived colon organoid and HT-29 cells. (A) Representative fluorescence microscopy images showing AUF1 expression in scramble morpholino treated control biopsy (S-MO), IBD patient biopsy, and AUF1 morpholino treated control biopsy (AUF1-MO_ samples. Nuclei were counterstained with DAPI. DIC and merged images are shown. Scale bars, 100 μm. (B) Quantification of AUF1 fluorescence intensity in S-MO, IBD, and AUF1-MO groups. (C) Representative images of intestinal organoids from S-MO and AUF1-MO groups and IBD patient tissues, showing bright-field morphology and FITC-dextran fluorescence. Scale bars, 100 μm. (D) Quantification of intestinal barrier permeability based on FITC-dextran (4 kDa) fluorescence. (E) Transepithelial electrical resistance (TEER) measurements over time in S-MO, LPS, and AUF1-MO treated HT-28, colon carcinoma cell line. (F) Relative mRNA expression of Occludin and Claudin-2 in S-MO and AUF1-MO treated HT-29 cells. Data are presented as mean ± SEM. Statistical significance was determined as indicated in the individual panels; ns, not significant.

### AUF1 Knockdown in HT-29 Cells: Impact on Occludin and Claudin-2 mRNA turnover and translational control

#### AUF1 regulates epithelial barrier integrity in HT-29 cells through coordinated control of tight junction gene expression

To further investigate the mechanistic relationship between AUF1 and epithelial barrier function, we used the human colonic epithelial carcinoma cell line HT-29. HT-29 cells were treated with increasing concentrations of AUF1-MO for 48 h, resulting in a dose-dependent reduction in AUF1 protein expression. A 0.75 µM of AUF1-MO produced the most pronounced knockdown and was therefore used for subsequent experiments (**Fig. S1D, E**). We assessed epithelial barrier integrity by measuring transepithelial electrical resistance (TEER). AUF1-MO-treated cells (AUF1-MO cells) exhibited a progressive and significant reduction in TEER over time compared with S-MO-treated cells (S-MO cells) indicating impaired barrier function. LPS-treated monolayers which induces inflammatory cytokine production in the monolayer served as a positive control and similarly displayed a time-dependent decrease in TEER (**Fig. 2E**). Consistent with the organoid data, quantitative RT-PCR showed that AUF1 knockdown reduced occludin expression while increasing claudin 2 expression compared with control cells (**Fig. 2F**).

Together, these findings establish a functional association between AUF1 deficiency and epithelial barrier disruption. Loss of AUF1 is accompanied by reduced Occludin and increased Claudin-2 expression in both primary colonic organoids and HT-29 epithelial monolayers, phenocopying the alterations observed in IBD-derived organoids. Given previous evidence that AUF1 regulates the stability of various mRNAs, we next investigated whether AUF1 controls the post-transcriptional stability of Occludin and Claudin 2 transcripts in AUF1-deficient and control HT-29 cells.

#### Measurement of t_1/2_ of Occludin and Claudin-2 mRNA in control and AUF1 knockdown HT-29 cells

The mRNA decay kinetics were quantified by determining transcript half-lives (t½, expressed in minutes) after blocking nascent RNA synthesis with Actinomycin D in AUF1-MO and S-MO treated HT-29 cells. The t½ values for Occludin mRNA in AUF1-MO and S-MO cells were observed to be 19 ± 2.34 min and 42.5 ± 5.51 min, respectively. Conversely, Claudin-2 mRNA exhibited t½ values of 60 ± 7.11 min in AUF1-MO cells and 38.5 ± 6.27 min in S-MO cells. Thus, a reciprocal trend in the turnover of Occludin and Claudin-2 transcripts was observed in response to AUF1 knockdown. Importantly, the housekeeping transcript GAPDH maintained an invariant t½ irrespective of AUF1 knockdown, thereby serving as an internal control (**Fig. 3A-C**).

**Figure 3.**
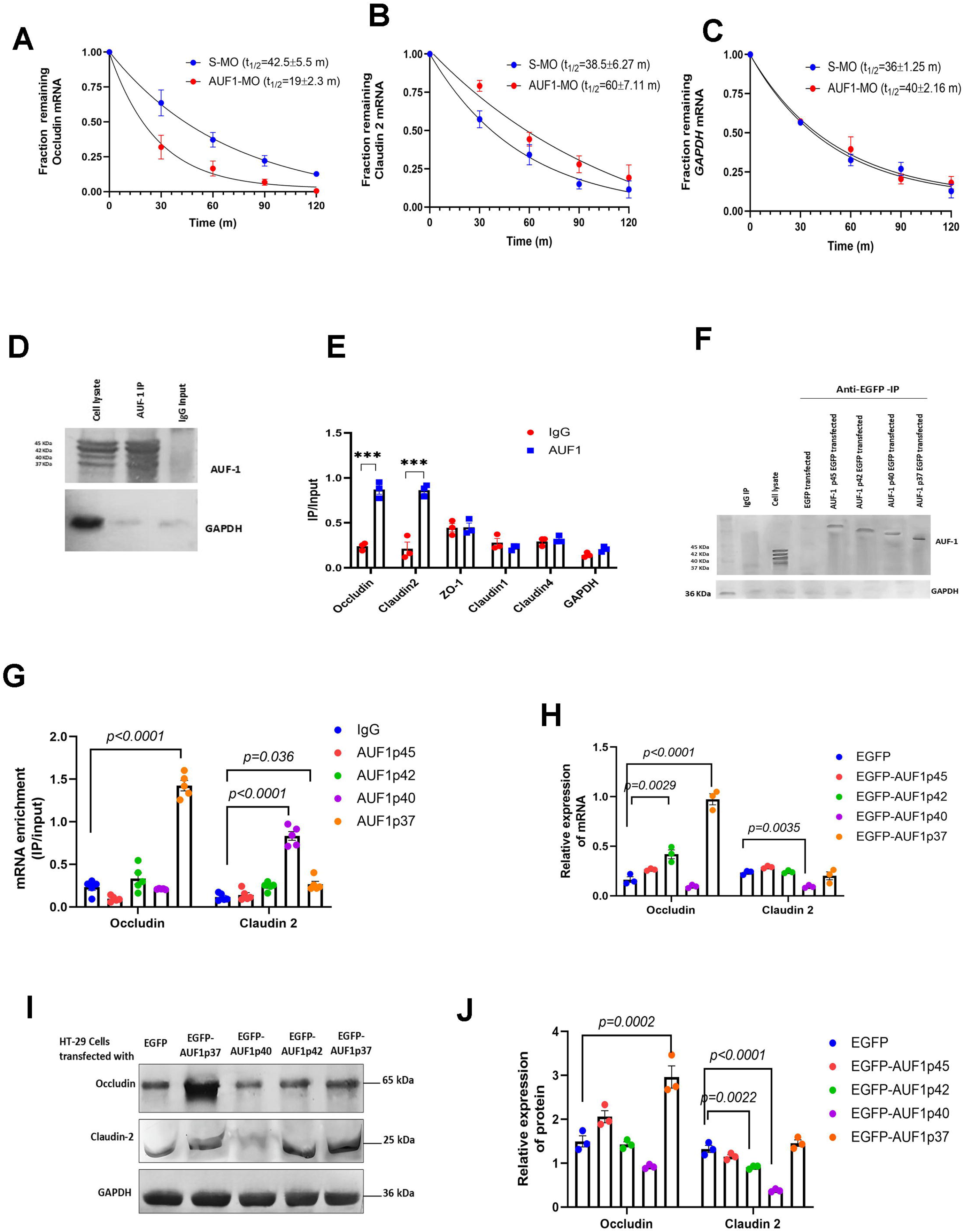
AUF1 regulates Occludin and Claudin-2 mRNA stability and expression. (A–C) mRNA decay kinetics showing the fraction of (A) Occludin, (B) Claudin-2, and (C) GAPDH mRNA remaining over time in control (S-MO) and AUF1-knockdown (AUF1-MO) HT-29 cells, including calculated mRNA half-lives. (D) Western blot analysis showing AUF1 immunoprecipitation (IP) compared with cell lysate and IgG controls, with GAPDH as a loading control. (E) Ribonucleoprotein immunoprecipitation (RIP) analysis showing the fold enrichment (IP/input) of target mRNAs (*Occludin, Claudin-2, ZO-1, Claudin-1, Claudin-4,* and *GAPDH*) associated with AUF1 relative to the control IgG. (F) Western blot confirmation of anti-EGFP immunoprecipitation in cells transfected with distinct EGFP-tagged AUF1 isoforms (p45, p42, p40, and p37). (G) Isoform-specific RIP assay measuring the enrichment (IP/input) of *Occludin* and *Claudin-2* mRNAs associated with individual AUF1 isoforms relative to control IgG. (H) Relative mRNA expression levels of *Occludin* and *Claudin-2* following overexpression of individual EGFP-AUF1 isoforms, compared with the EGFP control. (I) Western blot analysis of Occludin, Claudin-2, and GAPDH protein levels in HT-29 cells transfected with individual EGFP-AUF1 isoforms (p37, p40, p42, or p45) or EGFP control. (J) Quantification of relative Occludin and Claudin-2 protein expression levels normalized to the EGFP control from panel I. Data are presented as mean ± SD. Statistical significance was assessed using Student’s *t*-test or one-way ANOVA, as indicated by the specific *p*-values or asterisks.

#### Polysome profiling and translational status of Occludin and Claudin 2 mRNA

To interrogate the translational regulation of gene expression, polysome profiling was employed to evaluate the ribosomal engagement of Occludin and Claudin-2 mRNAs. Ribosome-associated transcripts from HT-29 cells were separated on a 0-50% continuous sucrose density gradient and analyzed by monitoring the gradient absorbance at 254 nm. This allowed to distinguish between polysomes, monosomes (80S or 40S/60S) and free RNA species. Thus, we found that in AUF1 silenced cells (AUF1-MO) Occludin mRNA was enriched in lighter fractions containing mainly monosomes (80S, 40S and 60S), whereas in S-MO cells Occludin mRNA was mostly localized in polysomes indicating its active translation. In contrast to Occludin, Claudin-2 mRNA in AUF1-MO cells was enriched in heavy polysome fractions, whereas in control cells it was distributed mainly in light fractions. As an internal control, GAPDH mRNA consistently localized to polysomal fractions irrespective of AUF1 status (**Fig. S2A-E**), thereby substantiating that AUF1 specifically orchestrates the translational regulation of Occludin and Claudin-2.

### Isoform-Dependent Interaction of AUF1 with Occludin and Claudin-2 mRNAs

#### Physical interaction of p37 and p40 with occludin and claudin-2 mRNA, respectively

To reveal physical association of Occludin and claudin-2 mRNA with AUF1, we performed an RNA immunoprecipitation (RNA-IP) assay using an anti-AUF1 antibody. The western blot showed the presence of AUF1 in anti-AUF1 IP, but not in control IgG, and qRT-PCR analysis showed the presence of Occludin and claudin-2 mRNA in anti-AUF1-IP and not in IgG-IP **(Fig. 3D, E)**.

With an assertion to identify AUF1 isoforms that may interact with occludin and claudin-2 mRNA, we transfected HT29 cells with or without plasmids either encoding EGFP or EGFP-AUF1p45 or EGFP-AUF1p42 or EGFP-AUF1p40, or EGFP-AUF1p37. RNA-IP was performed for each sample using anti-EGFP antibody. The western blot showed the presence of AUF1-GFP in anti-EGFP IP but not in control IgG IP. RNA-IP qRT-PCR analysis revealed that occludin and claudin-2 mRNAs preferentially associated with the p37 and p40 isoforms, respectively, whereas the other AUF1 isoforms showed no appreciable association, apart from a weak interaction between p37 and claudin-2 mRNA (**Fig. 3F, G**).

#### p37 upregulates Occludin expression, whereas p40 downregulates Claudin-2 mRNA expression

To ascertain whether the binding of AUF1 isoforms p37 and p40 influences the expression of Occludin and Claudin-2, respectively, we assessed TJ protein levels in HT-29 cells transfected with plasmids encoding EGFP-AUF1 isoforms. Both Western blot and qPCR were employed for this analysis. Isoform-specific qPCR primers confirmed that the transfected AUF1 isoforms were expressed at markedly elevated levels compared to their endogenous basal expression indicating the cells were optimally overexpressed with indicated AUF1 isoform. It was observed that transfection with EGFP-AUF1p37 markedly enhanced Occludin expression relative to other isoforms. Conversely, EGFP-AUF1p40 transfection produced a pronounced reduction in Claudin-2 expression, as corroborated by both qPCR and western blot analyses. Significantly, EGFP-AUF1p42 demonstrated relatively slight effects, resulting in a minor elevation of Occludin mRNA levels and a negligible decrease in Claudin-2 protein expression. (**Fig. 3H-J).**

Collectively, the data presented here demonstrate that AUF1 isoforms p37 and p40 associate with Occludin and Claudin-2 mRNAs, respectively, and regulate the expression of these proteins at the translational level. Thus, it is essential to determine the mechanisms by which AUF1 controls the stability of Occludin and Claudin-2 mRNAs.

### AUF1-miR122 Crosstalk Directing Occludin mRNA Stability and Counteracting miR-Induced Translational Repression

Earlier publications indicate that AUF1 has the ability to bind microRNAs [14], and that RNA-binding proteins like HuR can act in trans to protect their target mRNAs from microRNA-mediated repression, increasing their protein output [15]. Therefore, we decided to test whether microRNAs targeting Occludin and Claudin-2 could also bind AUF1. Indeed, miR122 regulates Occludin mRNA [16] and miR195 regulates Claudin-2 mRNA [17]. In this respect, to determine whether miR-122 could bind to AUF1 or miR-195 could bind to AUF1, HT-29 cells were transiently transfected with plasmids encoding miR122 or miR195. Interestingly, transfection of miR122 decreased Occludin protein levels in a dose-dependent manner, while Claudin-2 levels were not altered **(Fig. 4A).** As expected, increasing amounts of miR195 transfection decreased Claudin-2 protein levels, while Occludin levels were not changed **(Fig. 4B).**

**Figure 4.**
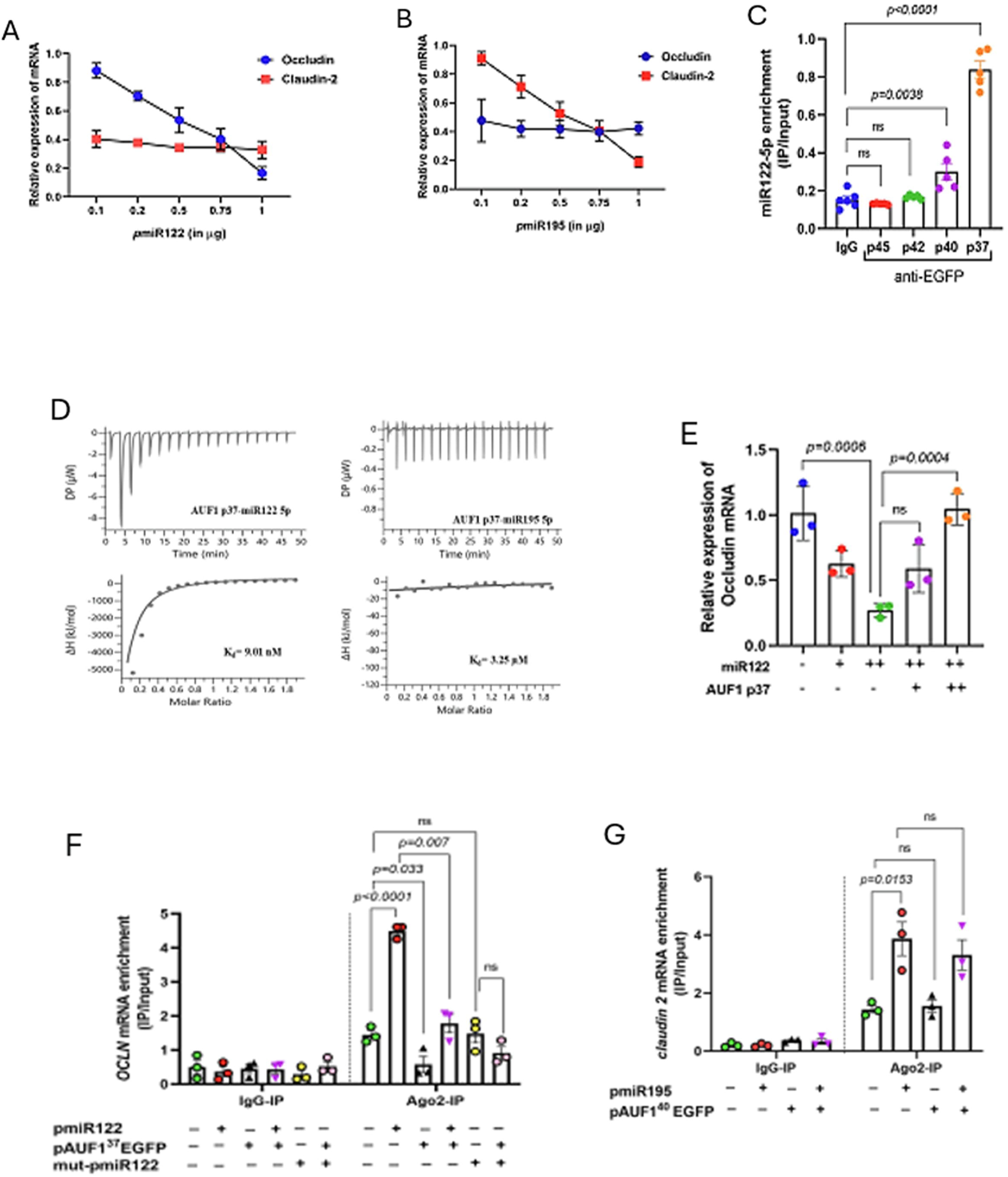
AUF1 Isoforms Differentially Regulate miR-122- and miR-195-Mediated Suppression of Occludin and Claudin-2. (**A, B**) Dose-dependent effects of miRNA expression on target mRNA levels. Relative mRNA expression levels of Occludin and Claudin-2) were measured following transfection with increasing concentrations (0.1–1.0 μg) of (A) pmiR122 or (B) pmiR195. (C) Interaction of miR-122-5p with AUF1 isoforms. RNA immunoprecipitation (RIP) assays were performed using anti-EGFP antibodies following tansfection of EGFP-tagged AUF1 isoforms (p45, p42, p40, or p37) in HT 29 cells or control IgG. miR-122-5p enrichment was quantified relative to input. (D) Isothermal titration calorimetry (ITC) analysis of AUF1 p37 binding affinity. Representative thermograms (top) and binding isotherms (bottom) show direct binding of AUF1 p37 to miR-122-5p (*K*d = 9.81 nM; left) and miR-195-5p (*K*d = 3.35 μM; right). (E) Rescue of miR-122-mediated Occludin repression by AUF1 p37. Relative Occludin mRNA expression was measured in cells transfected with increasing doses of miR-122 alone or in combination with AUF1 p37. (F, G) Ago2-mediated mRNA recruitment assessed by RIP-qPCR. (F) Enrichment of Occludin mRNA (*OCLN*) in IgG or Ago2 immunoprecipitates following transfection with pmiR122, pAUF137-EGFP, or mutant pmiR122 (mut-pmiR122), as indicated. (G) Enrichment of Claudin-2 mRNA in IgG or Ago2 immunoprecipitates following co-transfection of pmiR195 and pAUF1p40-EGFP. Data are presented as mean ± SD. Statistical significance was determined using one-way ANOVA or Student’s t-test, as appropriate (ns, not significant).

#### miR122, but not miR195, binds to AUF1 independently of its presence

RNA immunoprecipitation was performed with anti-EGFP antibody after transient transfection of cells with control EGFP plasmid or EGFP fused to AUF1 (p45, p42, p40, or p37). Subsequent analysis of microRNA enrichment within the immunoprecipitated complexes revealed that neither miR-122-3p (**Fig. S3A**) nor miR-195 (**Fig. S3B**) exhibited detectable association with any AUF1 isoform. In contrast, miR-122-5p was predominantly enriched in the p37 isoform immunoprecipitates, with a lesser but discernible association observed for the p40 isoform immunoprecipitates (**Fig. 4C**).

#### AUF1 binds miR122 but not miR195 with high affinity

RNA-protein interaction studies using isothermal titration calorimetry and a filter-binding assay showed an interaction between miR-122 and the AUF1p37 isoform, whereas there was no significant interaction with miR-195. This suggested that AUF1 could bind specifically to miR-122 RNA via the non-canonical ARE. The affinity of AUF1 for miR-122 was found to be in the nanomolar range, with a Kd of 9.01 nM, suggesting a significantly strong interaction between AUF1 and miR-122. **(Fig. 4D, E).**

### AUF1 Antagonizes miR-122–Mediated Translational Repression of Occludin mRNA

#### AUF1p37 displaces miR-122 from the Occludin mRNA binding site

We next investigated the functional consequence of AUF1p37 on miR-122–mediated translational repression of Occludin. AUF1 is known to impede miRNA biogenesis or maturation by promoting the decay of Dicer1 mRNA, which encodes the RNase III enzyme responsible for converting precursor miRNAs into their mature forms. To circumvent this confounding variable, all overexpression experiments employed plasmids encoding the mature sequences of miR-122 and miR-195. Consistent with prior reports, ectopic overexpression of miR-122 in HT-29 cells elicited a dose-dependent suppression of Occludin expression. Strikingly, co-expression of AUF1p37, that too in a high concentration, rescued Occludin expression even in the presence of the highest concentration of miR-122, thereby reversing the inhibitory effect (**Fig. 4F**). Importantly, AUF1p37 overexpression did not alter cellular miR-122 abundance (**Fig. S3C**), indicating that its antagonistic action was not mediated through miRNA degradation. Moreover, AUF1p37 expression had no discernible effect on the steady-state levels of Argonaute 2 (Ago2) (**Fig. S3D, E**), the principal effector of the RNA-induced silencing complex (RISC), suggesting that the observed reversal of repression was not attributable to RISC inactivation.

#### AUF1p37 prevents occludin mRNA from being associated with RISC

We next examined whether AUF1 influences the association of occludin and claudin-2 mRNAs with the RISC complex. RNA immunoprecipitation with anti-Ago2 antibody of cell lysates expressing miR-122 showed that occludin mRNA was bound to Ago2. However, AUF1p37 overexpression significantly reduced the association between occludin mRNA and Ago2, indicating that AUF1 facilitates the dissociation of occludin mRNA from the RISC complex. Consistently, AUF1p37 overexpression markedly reduced the association of miR-122 with Ago2, suggesting that AUF1p37 disrupts the interaction of miR-122 with both Ago2 and occludin mRNA. Supporting this, a mutant form of miR-122—harbouring a substitution in the occludin-binding region—failed to promote association of occludin mRNA with Ago2, regardless of AUF1p37 expression (**Fig. 4G**).

A parallel experiment was conducted to assess claudin-2 mRNA interaction with the RISC complex. In cells overexpressing miR-195, claudin-2 mRNA consistently associated with Ago2 irrespective of AUF1p40 overexpression (**Fig. 4H**). This indicates that AUF1p40 does not disrupt miR-195 binding to claudin-2 mRNA, and thus plays no role in its dissociation from the RISC complex.

### Coupled decay of AUF1 and Claudin-2 mRNA through ubiquitination

#### Proteasome is involved in the decay of claudin-2 mRNA

Earlier investigations have implicated AUF1 in promoting ARE–mRNA degradation via a proteasome-dependent pathway, potentially mediated through ubiquitination of specific AUF1 isoforms [18]. To assess whether such a mechanism underlies Claudin-2 decay, we overexpressed AUF1p37 or AUF1p40 in HT-29 cells and subjected them to treatment with or without the pan-proteasome inhibitor MG132. As anticipated, Claudin-2 expression was markedly reduced by AUF1p40 overexpression; however, this suppression was abrogated upon MG132 treatment, restoring Claudin-2 levels. In contrast, Occludin expression remained persistently elevated under AUF1p37 overexpression, irrespective of proteasome inhibition (**Fig. 5A**).

**Figure 5.**
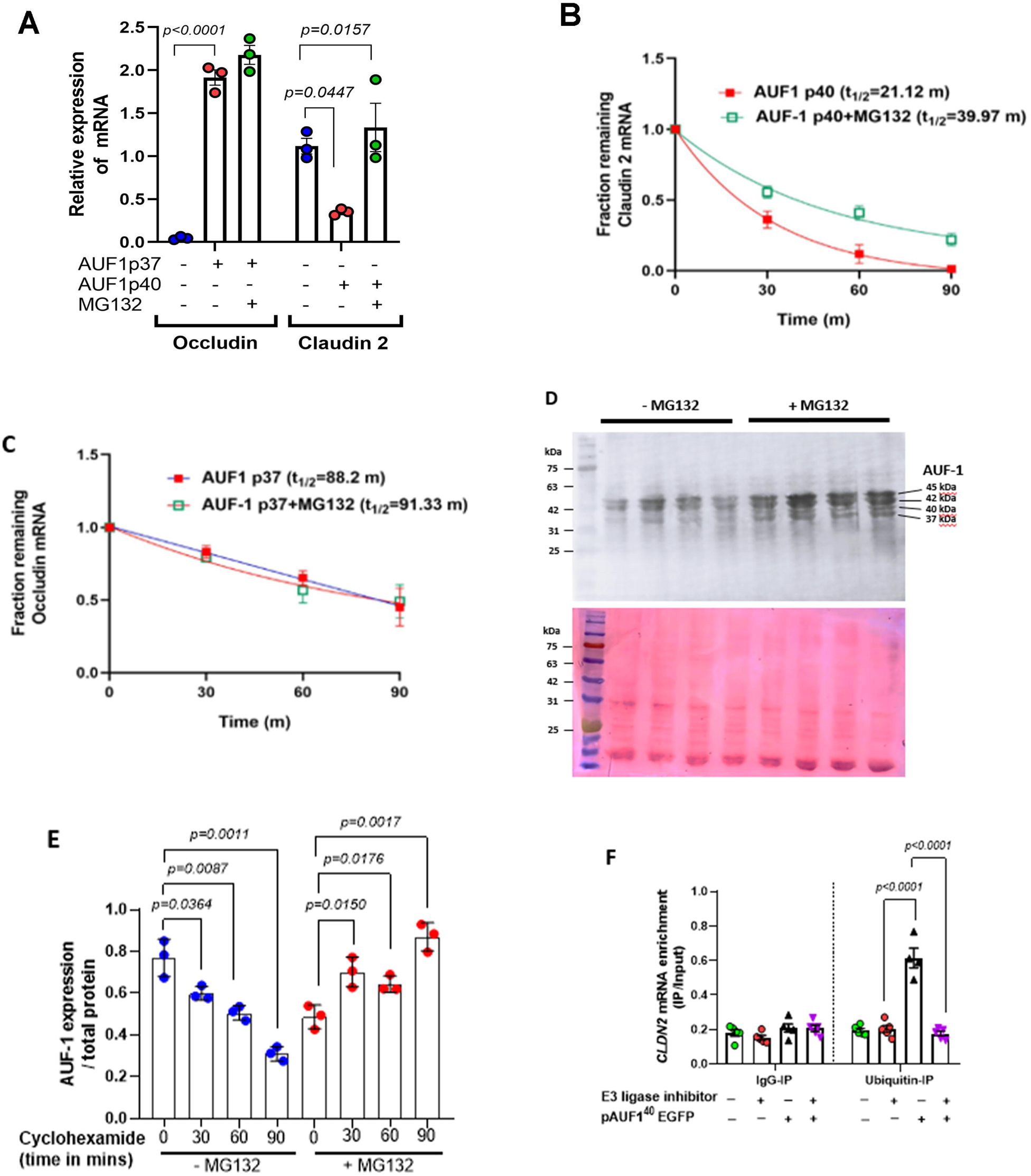
Proteasomal Inhibition Restores Claudin-2 mRNA Stability and AUF1 Protein Turnover. **(A)** Relative expression levels of Occludin and Claudin-2 mRNAs following transfection with AUF1 p37 or AUF1 p40 in the presence or absence of the proteasome inhibitor MG132. MG132 treatment rescued AUF1 p40-mediated suppression of Claudin-2 mRNA (*p* = 0.0157), whereas Occludin mRNA levels remained elevated (*p* < 0.0001). (B) mRNA decay kinetics of Claudin-2 in the presence of AUF1 p40 without MG132 (*t*_1/2_ = 21.12 min; red line) or with MG132 (*t*_1/2_= 39.97 min; green line) over a 90-min time course. (C) mRNA decay kinetics of Occludin in the presence of AUF1 p37 without MG132 (*t*_1/2_= 88.2 min; red line) or with MG132 (*t*_1/2_= 91.33 min; green line) over a 90-min time course. (D) Representative western blot analysis (top) and corresponding membrane/loading stain (bottom) showing AUF1 isoform levels (p45, p42, p40, and p37) over time under vehicle (-MG132) or proteasome inhibition (MG132) conditions. (E) Quantification of total AUF1 protein expression normalized to total protein levels following cycloheximide treatment at 0, 30, 60, and 90 min in the presence (+MG132) or absence (-MG132) of proteasome inhibition. (F) RNA immunoprecipitation (RIP) analysis showing enrichment of Claudin-2 mRNA (*CLDN2*) in IgG or ubiquitin immunoprecipitates under the indicated treatment conditions, with or without E3 ligase inhibitor and pAUF1p40-EGFP overexpression (*p* < 0.0001). Data are presented as mean ± SE. Statistical significance was determined using one-way ANOVA or Student’s t-test, as appropriate.

To further delineate this mechanism, transcript stability assays were performed to determine mRNA half-lives under these conditions. The half-life of Occludin mRNA was unaltered by MG132, whereas Claudin-2 mRNA showed a substantially prolonged half-life in the presence of the inhibitor **(Fig. 5B, C).** These results demonstrate that AUF1p40-mediated decay of Claudin-2 mRNA is proteasome-dependent, whereas AUF1p37-directed stabilization of Occludin mRNA is proteasome-independent.

#### AUF1 is degraded by ubiquitination

To determine AUF1 isoforms’ degradation kinetics, cells were treated with a translational inhibitor cycloheximide (CHX), and AUF1 stability was assessed by western blotting. Degradation of AUF1 was linear after CHX treatment with a calculated half-life of 76.87 min. At the same time, MG132 treatment prevented AUF1 degradation, and AUF1 protein levels were significantly higher than in the control cells at 30, 60, and 90 min after treatment, indicating that AUF1 is mainly subjected to ubiquitin-proteasome degradation pathway **(Fig. 5D, E)**.

#### Claudin-2 degraded with AUF1p40 through the process of ubiquitination

Next, we asked whether AUF1’s interaction with Claudin-2 and Occludin transcripts required ubiquitination. To test this, we used the E3 ubiquitin ligase inhibitor thalidomide that prevents the addition of ubiquitin chains to target proteins and transcripts. We performed qRT-PCR for Claudin-2 mRNA on immunoprecipitates obtained with anti-ubiquitin antibodies. This was done in cells overexpressing AUF1p40 with or without inhibitor treatment. AUF1p40 overexpression strongly increased the amounts of Claudin-2 mRNA in ubiquitin immunoprecipitates. However, this interaction was completely disrupted in cells treated with the E3 ligase inhibitor thalidomide. On the contrary, AUF1p37 overexpression, even in the presence of thalidomide, did not increase the levels of Occludin mRNA in ubiquitin immunoprecipitates **(Fig. 5F).**

### Generation and characterization of AUF1-engineered colonic organoids for regenerative therapy

Having established the role of AUF1 in maintaining epithelial barrier integrity by fine-tuning Occludin and Claudin-2, we asked whether restoring AUF1 expression could be beneficial in terms of epithelial repair. To this end, we generated colonic organoids from control and DSS-treated mice and engineered them to express EGFP-tagged AUF1p37 and p40. Plasmids encoding EGFP-AUF1p37 and EGFP-AUF1p40 protein were transfected into the organoids, and the stable expression of EGFP-AUF1 was validated 20 days after transfection. We first evaluated the phenotype of AUF1-engineered organoids derived from healthy control mice. These organoids exhibited inherent AUF1 isoforms at baseline, and the introduction of EGFP-AUF1p37 and EGFP-AUF1p40 led to a significant elevation in AUF1 levels. Notably, AUF1 overexpression in control-derived organoids was accompanied by increased Ki67 staining, indicative of enhanced proliferative activity **(Fig S4)**. Because excessive proliferation could represent an undesirable property for a cell-based therapeutic product, we next evaluated AUF1-engineered organoids generated from DSS-injured colonic tissue.

Organoids derived from DSS-injured mice similarly showed robust expression of EGFP-AUF1p37 and EGFP-AUF1p40 following transfection, but did not exhibit the increase in Ki67-positive cells observed in AUF1-engineered control-derived organoids **(Fig. S4)**. Thus, although AUF1 expression was effectively increased in both organoid populations, organoids derived from DSS-injured tissue displayed a more favourable functional phenotype for transplantation, with restoration of epithelial characteristics without evidence of excessive proliferation. We therefore selected AUF1-engineered organoids derived from DSS-injured tissue for subsequent regenerative transplantation experiments.

### AUF1-engineered colonic organoids promote epithelial regeneration and restore mucosal integrity in experimental colitis

We next evaluated whether transplantation of AUF1-engineered organoids could enhance tissue repair in vivo. AUF1 p37- and p40-expressing organoids derived from DSS-injured colons were transplanted by rectal delivery into mice with DSS-induced colitis **(Fig. 6A).** This approach was designed to determine whether restoring the AUF1-dependent epithelial program within transplanted cells could improve the regenerative capacity of organoid therapy in an inflamed and structurally damaged colon.

**Figure 6.**
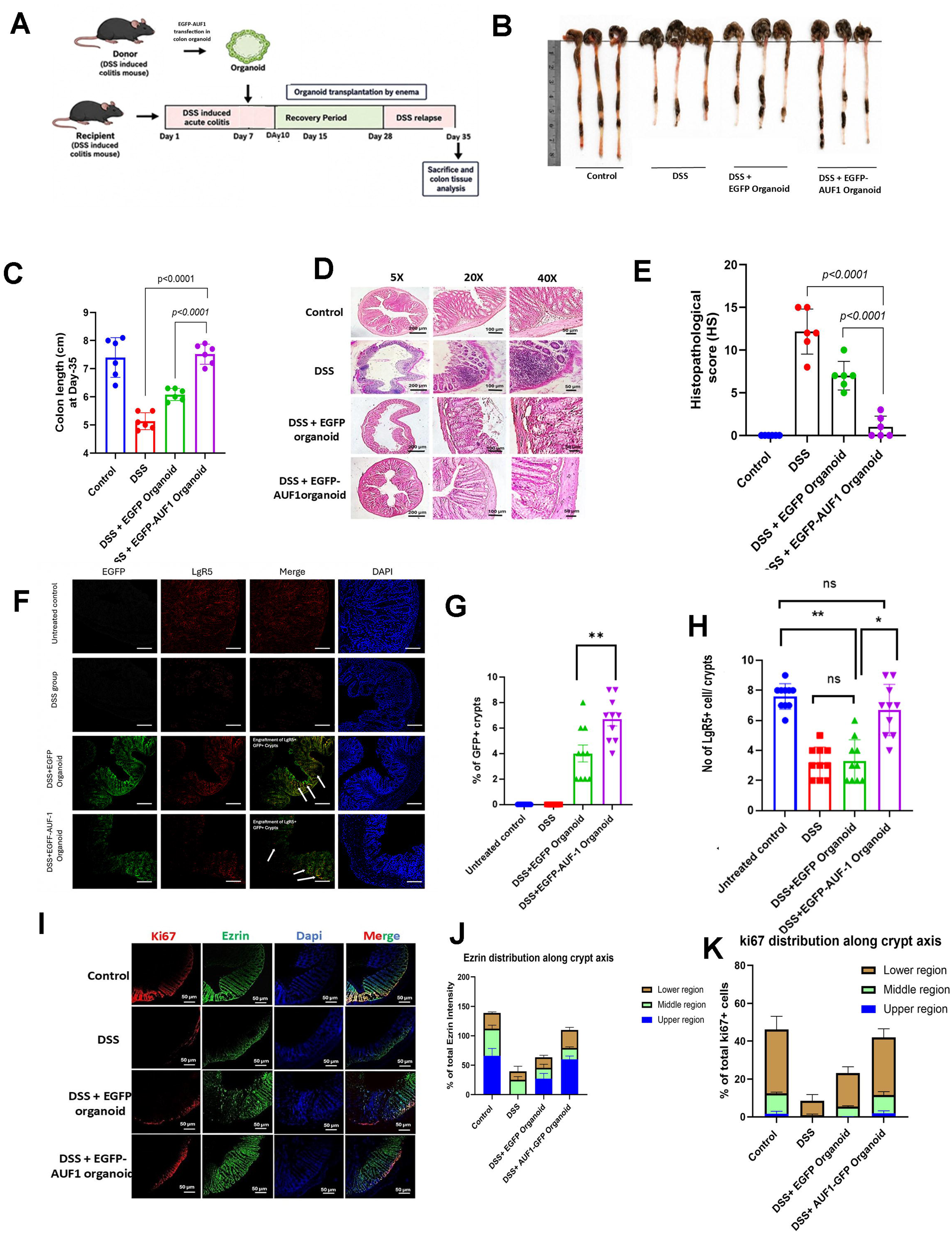
Transplantation of AUF1-Expressing Organoids Ameliorates DSS-Induced Colitis and Enhances Epithelial Regeneration. (A) Schematic representation of the experimental timeline for organoid isolation from donor mice, transfection/transduction, and enema-based transplantation into recipient mice with DSS-induced acute colitis. Mice underwent recovery and relapse phases before sacrifice on Day 35. (B, C) Gross examination and quantification of colon length. (B) Representative images of harvested colons and (C) quantitative analysis of colon length (cm) on Day 35 in control, DSS, DSS + EGFP organoid, and DSS + EGFP-AUF1 organoid groups (*p* < 0.0001). (D, E) Histopathological assessment of colon tissue. (D) Representative hematoxylin and eosin (H&E)-stained colon cross-sections at 5×, 20×, and 40× magnifications. (E) Histopathological scores (HS) across treatment groups (*p* < 0.0001). (F–H) Engraftment of transplanted organoids and expression of stem cell markers. (F) Representative immunofluorescence images showing EGFP (green), Lgr5 (red), merged signals (white arrows indicate co-localization), and DAPI nuclear counterstaining (blue) in colonic crypts. (G) Percentage of EGFP-positive (EGFP+) crypts following transplantation (*p* < 0.01). (H) Quantification of Lgr5+ stem cells per crypt across experimental groups (*p* < 0.01; *p* < 0.05; *ns*, not significant). (I–K) Epithelial proliferation and differentiation along the crypt axis. (I) Representative immunofluorescence images showing Ki67 (red; proliferation marker), Ezrin (green; apical membrane marker), DAPI (blue), and merged signals in the colonic epithelium. (J) Quantification of Ezrin fluorescence intensity across the lower, middle, and upper crypt regions. (K) Distribution of Ki67+ proliferating cells across the lower, middle, and upper crypt regions. Data are presented as mean ± SE. Statistical significance was evaluated using one-way ANOVA or Student’s t-test, as appropriate.

Following transplantation, in contrast to control EGFP-expressing organoids, AUF1-engineered organoids were associated with more extensive restoration of colonic architecture. Mice receiving AUF1-engineered organoids showed greater recovery of colon length, a macroscopic indicator of disease-associated tissue injury and recovery, compared with animals receiving control EGFP-organoids **(Fig. 6B, C)**. Histological analysis provided further evidence of enhanced mucosal repair **(Fig. 6D, E)**. Consistent with these results, AUF1-engineered organoids engrafted within areas of epithelial injury and were detected within damaged colonic crypts. The transplanted organoids progressively expanded during the recovery phase, consistent with their persistence and integration into the regenerating epithelium (**Fig. 6F-H**)

Additionally, AUF1-engineered organoid transplantation promoted restoration of crypt architecture and epithelial organization, with a corresponding increase in Ki67-positive epithelial cells, indicative of enhanced epithelial proliferative activity. Recovery of apical Ezrin expression further supported restoration of epithelial polarity and differentiation **(Fig. 6I-K)**.

Because persistent epithelial injury can promote pathological tissue remodelling, we next examined markers of fibrosis. AUF1-engineered organoid transplantation was associated with reduced accumulation of α-smooth muscle actin (α-SMA) and collagen III within the injured colon **(Fig. S5A–C).** These changes suggest that restoration of AUF1-dependent epithelial function not only promotes epithelial regeneration but may also limit the pathological remodelling that accompanies unresolved mucosal injury.

Together, these findings demonstrate that engineering intestinal organoids to restore AUF1 enhances their regenerative activity following transplantation into the injured colon. The improved epithelial architecture, proliferative response, epithelial polarity, and reduced fibrotic remodelling observed after AUF1-engineered organoid transplantation support the concept that restoring a disease-relevant epithelial regulatory program can improve the functional outcome of organoid-based therapy in experimental colitis.

## Discussion

Our findings identify AUF1 as a post-transcriptional regulator of intestinal epithelial barrier integrity and provide a mechanistic rationale for restoring AUF1 activity as part of a regenerative approach to IBD. Across human tissue, primary colonic organoids, epithelial monolayers, and experimental colitis, reduced AUF1 was associated with a characteristic TJ imbalance—loss of Occludin and induction of Claudin-2—and with impaired barrier function. AUF1 has previously been implicated in coordinated regulation of gene expression programs in multiple physiological contexts [12]; [19], and our findings extend this function to epithelial barrier maintenance. Mechanistically, AUF1 acted through isoform-specific regulation of these transcripts: p37 stabilized and promoted translation of Occludin, in part by antagonizing miR-122–RISC engagement, whereas p40 promoted Claudin 2 mRNA decay through a ubiquitin–proteasome–dependent pathway.

The translational value of these findings was validated by the concordance between human and model systems. Reduced epithelial AUF1 was observed in UC tissue and inversely correlated with the UCEIS. AUF1 reduction recapitulated the barrier dysfunction phenotype in human organoids similar to those derived from IBD patients. Our decision to focus on Occludin and Claudin-2 was based on their established and opposing roles in regulating epithelial permeability, with the former promoting barrier integrity [20] and the latter associated with enhanced cation selectivity and increased expression in inflamed intestine [9]; [21]. Consistent with previous observations in IBD, reduced Occludin and increased Claudin-2 were associated with disease activity [10]; [11]. These observations support a model in which loss of AUF1 contributes to, rather than merely reflects, epithelial dysfunction. Nevertheless, the human data are associative, and longitudinal studies will be required to determine whether AUF1 loss precedes disease activity or is predominantly induced by the inflammatory environment.

Our data further establish that AUF1 is not a uniform regulator of ARE-containing transcripts. Instead, its effects depend on isoform, target RNA, and regulatory context. AUF1 exists as four alternatively spliced isoforms—p37, p40, p42, and p45—with distinct RNA-binding properties [12]; [22]. In our experiments, p37 preferentially associated with occludin mRNA and miR-122, limiting Ago2 association and thereby preserving Occludin expression. This mechanism is consistent with previous evidence that RNA-binding proteins can compete with microRNAs for access to target transcripts [23]; [24]; [25] and with reported interactions between AUF1 and microRNA pathways [14]. RNA binding protein-dependent remodelling of RNA structure can alter accessibility of microRNA recognition sites, as described for other RNA-binding proteins [26].

In contrast, AUF1 p40 associated with Claudin 2 mRNA and promoted its turnover through a ubiquitin–proteasome–dependent pathway. AUF1 has been linked to the human exosome and to ARE-mRNA decay [27], while earlier work demonstrated coupling of AUF1-dependent mRNA degradation to polyubiquitination and proteasomal activity [28]; [29]. Our findings extend this model by suggesting that AUF1 p40 can couple Claudin 2 mRNA to ubiquitin-dependent decay. The parallel effects of AUF1 depletion on Claudin 2 mRNA stability and protein abundance, together with the effects of proteasome and ubiquitination inhibition, support this interpretation. However, the E3 ligase responsible and the molecular composition of the RNA–protein complex remains to be defined.

The convergence of these pathways provides a mechanistic explanation for the reciprocal TJ changes associated with AUF1 loss. AUF1 p37 preserves Occludin through protection from miR-122– dependent RISC engagement, whereas AUF1 p40 restrains Claudin-2 through accelerated transcript turnover. Such isoform-specific regulation may allow a single RNA-binding protein to coordinate opposing components of epithelial permeability. Whether this regulatory architecture extends to additional TJ transcripts or other AUF1-associated microRNAs remains an important question.

Our in vivo findings provide an initial test of the therapeutic implications of this mechanism. Previous work using AUF1-deficient mice showed that loss of AUF1 produces intestinal phenotypes that resemble inflammatory and metabolic perturbations [18], supporting a broader role for AUF1 in intestinal homeostasis. In our experimental colitis model, transplantation of AUF1-engineered organoids enhanced epithelial regeneration, improved crypt architecture, increased epithelial proliferation, restored apical Ezrin expression, and reduced fibrotic remodeling. These effects are consistent with the emerging concept that intestinal organoids can provide a regenerative source of epithelium and may complement conventional anti-inflammatory therapies. Recent preclinical and clinical efforts have highlighted both the potential and the limitations of autologous intestinal tissue or organoid transplantation [30]; [31].

The therapeutic effect of AUF1-engineered organoids is particularly relevant because organoid transplantation without AUF1 may not fully restore the molecular programs required for durable barrier function. In our study, engineering the graft to restore AUF1 activity improved epithelial repair in experimental colitis, suggesting that functional modification of the transplanted epithelium may enhance the efficacy of regenerative cell therapy. Rectal delivery by enema also provides a route that is conceptually compatible with endoscopic administration approaches used in early organoid-transplantation studies (Clinical Trial ID: jRCTb032190207).

Several questions remain before translation to human therapy. The relative contribution of individual AUF1 isoforms to human intestinal regeneration requires further definition, as does the identity of the ubiquitin machinery coupling AUF1 p40 to claudin2 decay. The durability, safety, biodistribution, and engraftment of engineered organoids will also need to be evaluated in clinically relevant models. Because AUF1 participates in diverse cellular processes, including inflammation, metabolism, senescence, and oncogenic pathways [12], approaches that selectively restore its epithelial activity may be preferable to systemic manipulation. In addition, the present therapeutic experiments were performed in experimental colitis and therefore do not establish efficacy in human IBD or demonstrate that AUF1 restoration is sufficient to overcome the complex inflammatory and immune components of disease.

Together, our results establish a mechanistic link between AUF1-dependent post-transcriptional control and intestinal barrier integrity and provide proof of concept that restoration of AUF1 in transplanted organoids can enhance mucosal repair. By combining molecular reprogramming of the epithelial graft with tissue replacement, this approach may offer a framework for regenerative therapies that address both epithelial structure and function in IBD.

## MATERIALS AND METHODS

### Collection of colon biopsies and patient characteristics

A total of 21 patients with active ulcerative colitis (UC), with UC endoscopic index of severity (UCEIS) scores ranging from 2 to 7 [32], were enrolled in the study (Listed in Table-4). In addition, a total of 9 patients undergoing colonoscopy in whom intestinal disease was ruled out were included as controls (Listed in Table-4). The study was approved by the Ethics Committee of the National Institute for Research in Bacterial Infections (ICMR-NICED/IECBMHR/003/2023), and all procedures were performed in accordance with the approved protocol. Patients were informed about the study in advance and provided written informed consent for the collection of additional colon biopsies during routine diagnostic endoscopy. These biopsies were subsequently used for RNA and protein expression analyses. Based on the objective of the study, patients with mild and moderate disease activity were collectively classified as having active UC.

### Differential Gene Expression Analysis and Volcano Plot

RNA-sequencing/transcriptomic data obtained from colon biopsy samples of patients with active ulcerative colitis (UC) and non-IBD controls were analysed to identify differentially expressed genes between the two groups. Differential gene expression analysis was performed by comparing normalized gene-expression values between the UC and control groups. Genes were considered differentially expressed based on predefined thresholds (p<0.05) for statistical significance and magnitude of change. The results were visualized as a volcano plot, with the log2 fold change (log2FC) plotted on the x-axis and the −log10 adjusted *P* value plotted on the y-axis using GraphPad Prism8. Genes meeting the predefined significance criteria were highlighted to distinguish significantly upregulated and downregulated transcripts from nonsignificant genes.

### RNA Extraction, cDNA Synthesis, and Quantitative real-time PCR

Total RNA was extracted from colon biopsy specimens or HT-29 cells using TRIzol Reagent (Invitrogen) according to the manufacturer’s instructions. The concentration and purity of the isolated RNA were determined using a Nanodrop. Equal amounts of RNA from each sample were reverse-transcribed into complementary DNA (cDNA) using a commercially available PrimeScript 1st strand cDNA Synthesis Kit (Takara Bio, Cat No. 6110A) according to the manufacturer’s protocol. Quantitative real-time PCR (qRT-PCR) was performed using the synthesized cDNA and gene-specific primers (Listed in Table-1) with a commercially available TB Green Premix Ex Taq II (Tli RNase H Plus) Kit (Takara Bio, Cat No. RR820A) and quantified with StepOne Real Time PCR System (Applied Biosystems). The thermal cycling conditions were set according to the manufacturer’s recommendations. Relative gene expression was calculated using the comparative Ct (2^−ΔΔCt) method [33], and normalized to gene expression of respective control group with GAPDH used as the endogenous control. For microRNAs, a stable small nuclear RNA, U6 snRNA was used as the endogenous control. Each sample was analysed in technical replicates, and gene expression levels were normalized to the corresponding reference gene.

**Table 1:**
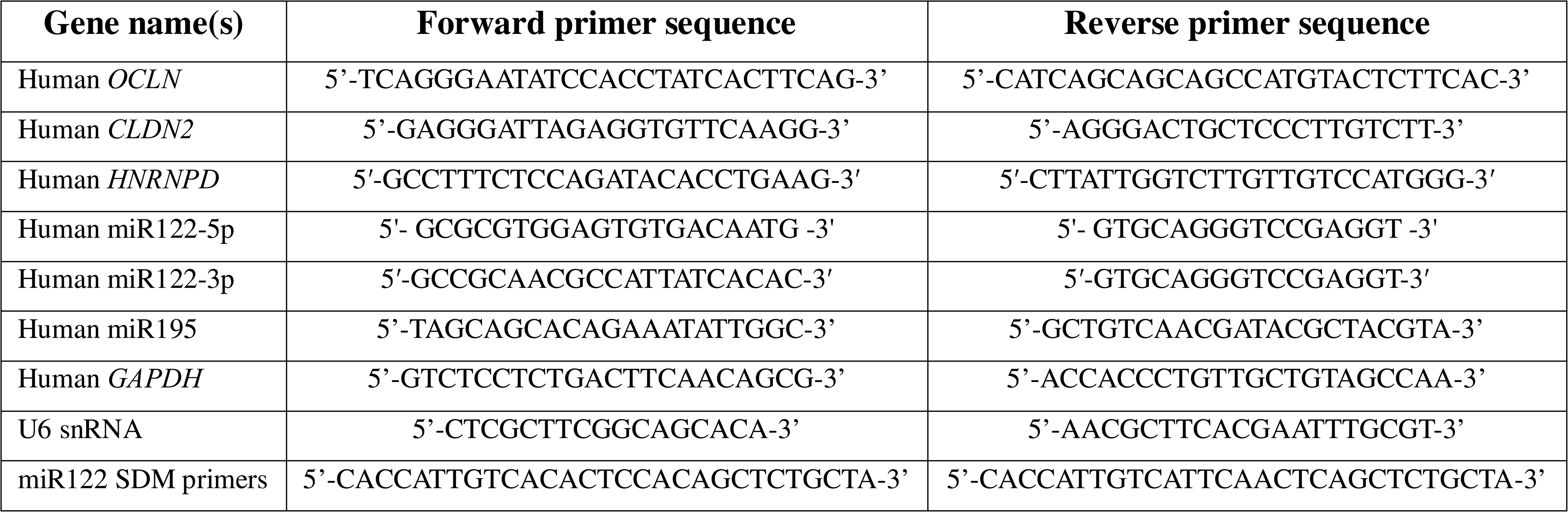
Primer sequences used for qRT-PCR and Site-directed Mutagenesis (SDM).

### Mice and animal ethics

C57BL/6 mice (6–8 weeks old) were obtained from the animal facility of ICMR-NICED, Kolkata, India. All animal experiments were approved by the Institutional Animal Ethics Committee of ICMR-NIRBI, Kolkata, India (PRO/2001/-Nov 2023-26), and were conducted in accordance with the guidelines of the Committee for the Purpose of Control and Supervision of Experiments on Animals (CPCSEA), Ministry of Environment and Forests, Government of India, New Delhi. Daily food intake was monitored by weighing the amount of food provided at the beginning of each 24-h period and the amount remaining at the end of the period. The difference between these measurements was taken as the daily food intake. Food consumption was recorded for each experimental group, with five mice per group.

### HT-29 Cell Culture

HT-29 human colorectal adenocarcinoma cells were maintained in RPMI-1640 medium (Gibco) supplemented with 10% Fetal Bovine Serum –US Origin (Corning) and 1% Penicillin–Streptomycin (Gibco). Cells were cultured at 37°C with 5% CO in a humidified incubator (Heracell 150i, ThermoFisher Scientific) and passaged upon reaching approximately 80–90% confluence. Cells were routinely examined for morphology and maintained under standard culture conditions until use in subsequent experiments.

### Plasmid constructs and isolation

For overexpression of AUF1 isoforms in HT-29 cells and organoids, four different pZeoSV2(+) plasmid constructs encoding GFP-tagged AUF1 isoforms, namely GFP-p45^AUF1^, GFP-p42^AUF1^, GFP-p40^AUF1^, and GFP-p37^AUF1^, were used. As subsequent GFP control, pEGFP-C1 (4.7 kb) plasmid was used. miR-122 was overexpressed using a plasmid construct containing the pre-miR-122 sequence cloned into the pTZ U6+1 vector, as described previously. Similarly, miR-195 overexpression was achieved using a plasmid construct containing the pre-miR-195 sequence cloned into the pSilencer 4.1 vector. For recombinant AUF1-p37 protein purification, a separate plasmid construct containing the cDNA encoding AUF1-p37 cloned into the pBAD/HisB vector was used. All plasmid constructs were transformed into chemically competent *Escherichia coli* DH5α cells using the CaCl_2_-mediated transformation method and subsequently propagated in Luria–Bertani (LB) broth supplemented with the appropriate selection antibiotic. Plasmid DNA was isolated using the SPINeasy Plasmid Miniprep Kit (MP Biomedicals) according to the manufacturer’s instructions.

### Site-directed mutagenesis (SDM)

Site-directed mutagenesis was performed to generate the desired substitution mutations in the pre-miR122 construct into the occludin mRNA-binding site. Briefly, mutation-specific primers containing the desired nucleotide substitutions were designed. 5 dis-continuous bases among the 8 bp occludin mRNA binding site of pre-miR122 (TGGAGTGT) was mutated, where the subsequent mutations were introduced in the reverse primer. Mutated pre-miR122 (Mut-pmiR122) construct were generated using Phusion Site-directed Mutagenesis Kit (ThermoFisher Scientific) according to manufacturer’s protocol. and subsequently transformed into competent *Escherichia coli* DH5α cells. Individual colonies were screened by plasmid isolation and running control experiments. The verified mutant plasmids were subsequently used for downstream expression and functional analyses.

### Cell transfection

For transfection, HT-29 cells were seeded at an appropriate density (approximately, 2 x 10^5^ cells for a 12-well plate) and allowed to adhere and grow up to 70-90% confluency before transfection. Plasmid DNAs were transiently transfected with Lipofectamine 2000 Transfection Reagent (Invitrogen) according to the manufacturer’s (Invitrogen) instructions. Briefly, plasmid DNA and Lipofectamine 2000 were separately diluted in serum-free medium, combined (1:3), and incubated for approximately 25 min at room temperature to allow complex formation. The complexes were then added dropwise to the cells and incubated under standard culture conditions. After an incubation period of 4-6 hours, the transfection medium was replaced with complete culture medium. Transfected HT-29 cells can be observed in next 36-72 h and were harvested at the indicated time points for subsequent molecular and functional analyses. Appropriate untreated and/or negative-control transfections were included in each experiment.

### In Vitro Knockdown of AUF-1 using AUF1 GMO-PMO

Human colon organoids and HT-29 cells were used for in vitro knockdown of AUF1 using a cell-penetrating gene-modulating oligomer (GMO)-phosphorodiamidate morpholino oligomer (PMO) targeting AUF1. Cells or organoids were treated with AUF1 GMO-PMO (AUF1-MO; Sequence in Table-2) at the indicated concentration for 48 hours. A sequence-matched control GMO-PMO (Scrambled-MO; Sequence in Table-2) was used as a negative control. Control and AUF1 GMO-PMO-treated samples were subsequently used for downstream functional analyses.

**Table 2:**
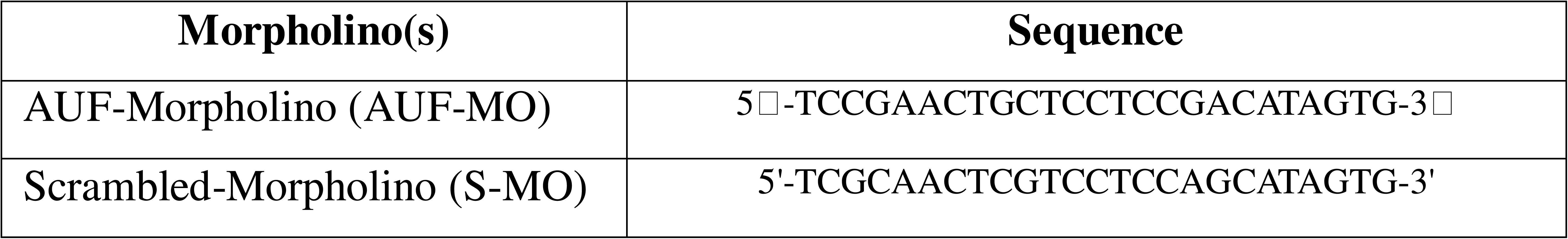
Morpholino-oligomer sequences used for gene knockdown.

### Development of Mouse and Human Colon Organoids

Colon organoids were established from mouse colon tissue or human colon biopsy specimens according to the protocols of STEMCELL Technologies. Briefly, human or mouse colon tissues were washed thoroughly with PBS (MP Biomedicals) supplemented with gentamicin (Gibco), minced into small fragments, and subjected to enzymatic dissociation using a Gentle Cell Dissociation Reagent (STEMCELL Technologies, Cat No. 100-0485) to release intestinal crypts. Isolated crypts were embedded in Matrigel Basement Membrane Matrix (Corning, Cat No. CLS354234) and cultured in specific IntestiCult Organoid Growth Medium – Human (STEMCELL Technologies, Cat No. 06010) or IntestiCult Organoid Growth Medium – Mouse (STEMCELL Technologies, Cat No. 06005) at 37°C in a humidified incubator with 5% CO. The medium was replaced every 2–3 days, and organoids were passaged upon reaching appropriate growth and density.

### Assessment of Epithelial Barrier Permeability

*FITC–Dextran Permeability Assay in Colon Organoids –* Colon organoids were incubated with Fluorescein Isothiocyanate-Dextran (FITC-Dextran; Sigma-Aldrich, Cat No. FD4) for the indicated time period. Following incubation, IN/OUT fluorescence intensity in the organoid culture medium was measured using a confocal microscope (Carl Zeiss, Germany). Epithelial permeability was determined based on the amount of FITC–dextran that crossed the organoid epithelial barrier, with higher fluorescence indicating increased permeability [34].

*TEER Measurement in HT-29 Cells –* HT-29 cells were seeded onto 12 well costar transwell cell culture inserts (Corning) and cultured until a confluent monolayer was established. Transepithelial electrical resistance (TEER) was measured using a Millicell ERS-2 Voltohmmeter (Millipore). Background resistance from cell-free inserts was subtracted from the measured values, and TEER was expressed as Ω·cm² after normalization to the surface area of the transwell membrane [35].

### mRNA Half-Life Analysis

To determine the mRNA decay kinetics mRNA half-life was quantified as described previously [36, 37]. Briefly, HT-29 cells were treated with actinomycin D (MP biomedicals) to inhibit de novo transcription, and cells were harvested at the indicated time points (0, 30, 60, 90, and 120 mins) following treatment. Total RNA was isolated from each sample, and the abundance of the target mRNA was quantified by qRT–qPCR. mRNA levels at each time point were normalized to the corresponding level at 0 h. The relative mRNA abundance was plotted against time, and the mRNA half-life was determined by fitting the decay data to a first-order exponential decay model.

### Polysome profiling

Polysome profiling was performed to assess the association of specific mRNAs with actively translating ribosomes, as previously described [38]. Briefly, cells were treated with 100 µg/mL cycloheximide (MP biomedicals) for approximately 30 mins to stabilize ribosome–mRNA complexes and lysed under polysome-preserving conditions. The clarified lysates were layered onto a linear sucrose density gradient (10-60%) and subjected to ultracentrifugation to separate free ribosomal subunits, monosomes, and polysomes according to their sedimentation properties. Gradients were fractionated while continuously monitoring absorbance at 254 nm to generate polysome profiles. RNA was isolated from individual fractions, followed by reverse transcription and quantitative real-time PCR to determine the distribution of target mRNAs across the fractions. The relative abundance of each transcript in polysomal versus non-polysomal fractions was used to assess its association with actively translating ribosomes.

### RNA immunoprecipitation

RNA immunoprecipitation (RIP) was performed to assess the association of target mRNAs with RNA-binding proteins, as previously described [39, 40] with minor modifications. Briefly, cells were lysed in an ice-cold RIP lysis buffer containing 40 mM Tris-HCl (pH −7.5), 150 mM NaCl, 0.5% NP-40 (ThermoFisher Scientific), 0.5% BSA supplemented with 40 U/mL RNase inhibitor (Applied Biosystems) and protease inhibitor cocktail (ThermoFisher Scientific). The lysates were pre-cleared with Protein G magnetic beads (Dynabeads Protein G for Immunoprecipitation, Invitrogen, Cat No. 10003D) and subsequently incubated overnight at 4°C with an antibody against the proteins of interest (Listed in Table-3) or an appropriate IgG mock antibody (Listed in Table-3) with gentle rotation. Protein G magnetic beads were then added and incubated for an additional period to capture the antibody–protein–RNA complexes. The beads were washed extensively with RIP wash buffer containing 40 mM Tris-HCl (pH −7.5), 150 mM NaCl, 0.5% NP-40 supplemented with 40 U/mL RNase inhibitor, and the immunoprecipitated RNA was subsequently isolated from the bead-bound complexes. Purified RNA was reverse-transcribed into cDNA, and the enrichment of target transcripts was quantified by quantitative real-time PCR. Relative enrichment was calculated by comparing the abundance of target mRNAs in the specific antibody immunoprecipitate with that in the corresponding mock IgG and/or input sample.

**Table 3:** Antibodies used for Western blotting, Immunofluorescence and Immunohistochemistry.

| Antibody name(s) | Host | Catalogue No. | Company |
| --- | --- | --- | --- |
| Occludin pAb | Rabbit | GTX114949 | GeneTex |
| Claudin-2 pAb | Rabbit | ab53032 | Abcam |
| AUF1 pAb (For Western blot) | Rabbit | 07-260 | Millipore |
| AUF1 pAb (For Immunofluorescence) | Rabbit | ab61193 | Abcam |
| Argonaute-2 (Ago2/EIF2C2) pAb | Rabbit | GTX131422 | GeneTex |
| GFP pAb | Rabbit | BB-AB0065 | Bio Bharati Life Sciences |
| Ki67 mAb | Rabbit | A20018 | ABclonal Technology |
| Ezrin (EPR23353-55) mAb | Rabbit | ab270442 | Abcam |
| α-SMA (ACTA2) mAb | Rabbit | A17910 | ABclonal Technology |
| Collagenase-III α1/COL3A1 pAb | Rabbit | A3795 | ABclonal Technology |
| GAPDH (14C10) mAb | Rabbit | 2118T | Cell Signalling Technologies |
| Lgr5/GPR49 mAb | Rabbit | A12327 | ABclonal Technology |
| Anti-Rabbit IgG (Alexa Fluor 488) | Goat | ab150077 | Abcam |
| Anti-Rabbit IgG (PE) | Goat | ab72465 | Abcam |
| Anti-Rabbit IgG (AP) | Goat | ab97048 | Abcam |
| Anti-Rabbit IgG (HRP) | Goat | ab205718 | Abcam |
| IgG Isotype Control | Rabbit | 31235 | Invitrogen |

**Table 4:** Clinical details of UC and healthy control patients from whom colon biopsy samples were collected.

| Patient ID | Age/Sex | UCIES Score | Colonoscopy finding(s) | Group |
| --- | --- | --- | --- | --- |
| S1 | 56 Y / F | 1+1+0= 2 | Rectal Ulcer | UC |
| S2 | 48 Y / F | 1+1+1= 3 | Rectal ulcer | UC |
| S3 | 22 Y / F | 1+0+2= 3 | Rectal ulcer | UC |
| S5 | 44 Y / F | 2+1+2= 5 | Patchy loss of vascularity, multiple large superficial and deep ulcer seen | UC |
| S6 | 50 Y / F | 2+1+2= 5 | Patchy loss of vascularity, multiple large superficial and deep ulcer seen | UC |
| S7 | 65 Y / M | N/A | Normal mucosa | Control |
| S8 | 24 Y / M | 2+1+2= 5 | Complete loss of vascular pattern, multiple superficial ulcers, coagulated blood seen | UC |
| S9 | 60 Y / M | N/A | Normal mucosa | Control |
| S10 | 59 Y / M | N/A | Normal mucosa | Control |
| S11 | 49 Y / F | N/A | Chronic malignancy with rectal ulcer | UC |
| S12 | 33 Y / M | 2+2+3= 7 | Continuous loss of vascular pattern with some spots and streaks of blood, superficial and deep ulcer | UC |
| S13 | 36 Y / M | N/A | Control mucosa | Control |
| S15 | 55 Y / F | 2+1+3= 6 | Rectal bleeding | UC |
| S17 | 28 Y / F | 1+0+1= 2 | Rectal bleeding | UC |
| S18 | 56 Y / M | 1+1+0= 2 | Sigmoid ulcer | UC |
| S19 | 64 Y / F | N/A | Normal mucosa | Control |
| S20 | 15 Y / M | 1+0+2= 3 | Rectal bleeding | UC |
| S21 | 59 Y / M | 1+1+1= 3 | Rectal bleeding | UC |
| S22 | 48 Y / F | N/A | Normal mucosa | Control |
| S24 | 52 Y / F | 1+1+0= 2 | Multiple ulcer size 1mm x 1mm to 3mm x 1mm seen, with erosions, no luminal bleed | UC |
| S25 | 24 Y / F | 2+2+2= 6 | Complete loss of vascular pattern, luminal bleed, superficial ulcer | UC |
| S26 | 38 Y / M | 2+0+2= 4 | Multiple ulcers, with erosions, luminal bleed | UC |
| S27 | 49 Y / F | 1+0+0= 2 | In remission | Control |
| S28 | 42 Y / F | N/A | Normal mucosa | Control |
| S29 | 23 Y / F | 1+2+0= 3 | Complete loss of vascularity with granularity, multiple erosion with no luminal bleed noted | UC |
| S30 | 36 Y / M | 1+1+1= 3 | Complete loss of vascularity with granularity, multiple erosion with no luminal bleed noted | UC |
| S31 | 35 Y / F | 2+0+0= 2 | Superficial ulcers, rest normal mucosa | UC |
| S32 | 16 Y / M | 2+2+2= 6 | Complete loss of vascular pattern, luminal bleed, superficial ulcers | UC |
| S33 | 39 Y / M | 2+2+0= 4 | Complete loss of vascular pattern, luminal bleed, superficial ulcers | UC |
| S37 | 37 Y / M | N/A | Normal mucosa | Control |

### Cycloheximide chase assay

Protein half-life was determined using a cycloheximide (CHX) chase assay to inhibit de novo protein synthesis, as previously described [41, 42] with minor modifications. Briefly, cells were treated with cycloheximide (100 µg/mL), and cells were harvested at t time points (0, 30, 60, and 90 mins) following treatment. At each time point, cells were lysed in ice-cold RIPA lysis buffer containing protease inhibitors, and equal amounts of total protein were subjected to SDS-PAGE followed by Western blotting. Band intensities were quantified by densitometric analysis and normalized to total protein loaded determined by ponceau staining of the same blot. Protein abundance at each time point was expressed relative to the level at 0 min.

### Western blotting

Cells or tissue samples were lysed in RIPA lysis buffer (Invitrogen) supplemented with protease inhibitor cocktail. Protein concentrations were determined using Pierce BCA Protein Assay Kits (ThermoFisher Scientific), and equal amounts of protein were mixed with Laemmli loading buffer (with β-mercaptoethanol) and denatured by heating. The protein samples were separated by SDS-PAGE and subsequently transferred onto PVDF membranes (Millipore). Membranes were blocked with 5% BSA in Tris-buffered saline (TBS; G-Biosciences) containing 0.1% Tween-20 (G-Biosciences) and incubated overnight at 4°C with the respective primary antibodies (Listed in Table-3) at a dilution of 1:1000. Following washing with TBST, membranes were incubated with specific AP-conjugated secondary antibody (Listed in Table-3) at a dilution of 1:10000 for 2 h at room temperature. After washing, protein bands were visualized using 1-Step NBT/BCIP Substrate Solution (ThermoFisher Scientific, Cat No. 34042); [43]. Band intensities were quantified by densitometric analysis and normalized to the corresponding loading control.

### Recombinant AUF1 protein expression and purification

Recombinant AUF1 protein was expressed and purified from bacterial cells as previously described [44] with few modifications. The pBAD/HisB expression constructs carrying p37^AUF1^ was transformed into *Escherichia coli* Rosetta2 (DE3) cells and recombinant His_6_-tagged AUF1 proteins were expressed following induction with 0.02% arabinose. The recombinant proteins were purified from bacterial lysates using Ni²-affinity chromatography with a PD-10 desalting column (Sigma-Aldrich). Briefly, induced bacterial cell pellets were harvested, lysed in Ni-NTA lysis buffer containing, 50mM Sodium Phosphate, pH 8.0, 500 mM NaCl, 20 mM imidazole (Sigma-Aldrich) and 1% Triton X-100 (Sigma-Aldrich) by sonication, and clarified by centrifugation. Following resin loading and washing with Ni-NTA lysis buffer was performed to improve protein purity before elution, which are thereafter eluted with a buffer containing high concentration of imidazole (300 mM). The eluted protein was subsequently dialyzed against 10 mM HEPES-KOH buffer (pH 7.5) using a dialysis membrane-110 (HiMedia) to remove excess imidazole and exchange the buffer. Recombinant protein purity was assessed by SDS-PAGE, and protein concentration was determined using BCA assay.

### Isothermal Titration Calorimetry (ITC)

The direct interaction between the purified p37^AUF1^ and microRNAs were evaluated by isothermal titration calorimetry (ITC) as previously described [45] with few modifications. Purified p37^AUF1^ and microRNAs were extensively dialyzed against the same HEPES-KOH buffer (pH −7.5) and degassed prior to analysis. The microRNA (2.5 µM) was loaded into the ITC sample cell, while the p37^AUF1^ protein (25 µM) was loaded into the injection syringe. Sequential injections of were performed into the microRNA solution at a constant temperature (25°C), and the heat released or absorbed during each injection was recorded using MicroCal PEAQ-ITC system (Malvern Panalytical, United Kingdom). A control titration of protein into assay buffer was performed to account for heat effects associated with dilution and injection. The corrected binding isotherm was analysed using a one-site binding model to determine the binding affinity (K_d_) with the help of MicroCal PEAQ-ITC Analysis Software (United Kingdom).

### Histological analysis

Colon tissues were collected from the experimental mice, gently flushed PBS, and fixed in 10% formalin (Sigma). The fixed tissues were processed, embedded in paraffin, and sectioned at approximately 4μm thickness using a regular microtome. Tissue sections were deparaffinized, rehydrated through graded ethanol, and stained with hematoxylin and eosin (H&E; Sigma-Aldrich) following standard protocols. The stained sections were examined under a light microscope for assessment of epithelial integrity, mucosal architecture, inflammatory cell infiltration, crypt damage, and ulceration. Histological changes were evaluated and scored using an established histopathological scoring system [46] by an investigator blinded to the experimental groups.

### Immunohistochemistry

Paraffin-embedded human colon biopsy tissue sections were cut into 4-μm-thick sections and mounted on positively charged glass slides (Sigma-Aldrich). Sections were deparaffinized in xylene (MP Biomedicals) and rehydrated through a graded ethanol series. Antigen retrieval was performed using an antigen-retrieval buffer under heat-induced conditions. Following cooling, sections were treated with hydrogen peroxide (Supelco, MERCK) to quench endogenous peroxidase activity and subsequently blocked to minimize nonspecific antibody binding.

Sections were incubated overnight at 4°C with the respective primary antibodies against proteins of interest (Listed in Table-3) at the optimized dilution (commonly, 1:200). After washing with PBS, sections were incubated with the appropriate HRP-conjugated secondary antibody. Immunoreactivity was visualized using a DAB liquid substrate system (Sigma-Aldrich), followed by counterstaining with hematoxylin. Sections were subsequently dehydrated, cleared, and mounted for microscopic examination. Images were acquired using a bright-field microscope under identical imaging conditions. Negative controls were included by omitting the primary antibody [47].

### Immunofluorescence staining

Paraffin-embedded human colon biopsy tissue sections or mice colon sections were cut into 4-μm-thick sections and mounted on positively charged glass slides. For immunofluorescence (IF) staining, sections were deparaffinized in xylene (MP Biomedicals) and rehydrated through a graded ethanol series. 50 µM ammonium chloride was used as quencher and after washing with PBS, samples were permeabilized and blocked with 5% FBS containing 3% BSA (Gold Bio) to minimize nonspecific antibody binding. Antigen retrieval was performed using citrate buffer (pH – 6) under heat-induced conditions. After washing in PBS, samples were then incubated overnight at 4°C with primary antibodies against the proteins of interest (Listed in Table-3) at an optimized dilution (commonly, 1:200). Following washing with PBS, the samples were incubated with appropriate fluorophore-conjugated secondary antibodies (1:200; Listed in Table-3) for 1 hour at room temperature in the dark. Nuclei were counterstained with DAPI (Sigma-Aldrich). After thorough washing, stained samples were mounted using DPX (Sigma-Aldrich) and examined using a confocal microscope (Carl Zeiss, Germany). Appropriate negative controls, including samples processed without primary antibody, were included to assess nonspecific staining. The fluorescence intensity was quantified using ZEN software (Gottingen, Germany) and expressed as corrected total cell fluorescence (CTCF). CTCF = Integrated density – (Area of selected cells x Mean fluorescence of background readings); [48].

### Generation of AUF1-engineered organoids for regeneration therapy

To generate AUF1-overexpressing organoids for regenerative therapy in mice, two different pZeoSV2(+) plasmid constructs encoding GFP-tagged AUF1-p37 (pEGFP-AUF1p37) or AUF1-p40 (pEGFP-AUF1p40) were co-transfected into DSS-derived organoids as described previously [49], with modifications. Briefly, organoids were dissociated into single cells or small cell clusters using the gentle cell dissociation reagent and subsequently transfected with the respective cloned pZeoSV2(+) plasmids (containing a Zeocin resistance cassette) carrying pEGFP-AUF1p37 or pEGFP-AUF1p40.

Following transfection, the cells were embedded in Matrigel and allowed to recover before selection was initiated using organoid culture medium containing 25 µg/mL Zeocin (Invitrogen). Surviving organoids were expanded and serially passaged under continuous Zeocin selection to enrich for EGFP-AUF1p37 and pEGFP-AUF1p40 co-expressing organoids. Stable expression of the AUF1 constructs was monitored for 20 days following transfection, and expression was confirmed by EGFP fluorescence using confocal microscopy before their use in subsequent regenerative therapy experiments.

### Induction of DSS-Induced intestinal injury

DSS-induced intestinal injury was established in C57BL/6 mice (6–8 weeks old male) by administering 2.5% (w/v) dextran sodium sulphate (DSS; molecular mass, 36–50 kDa) dissolved in purified water to mice for 7 days [50]. 28 days after rectal administration of organoids, 2.5% DSS was again reintroduced via drinking water for 7 days (DSS-relapse). After second phase of DSS administration, following euthanasia, the colon was carefully excised from the ileocecal junction to the anus and gently laid out without stretching. The colon length was measured using a ruler, and the measurement was recorded in centimetres (cm). Colon shortening was assessed as an indicator of DSS-induced intestinal injury.

### Rectal administration of AUF1-engineered organoids

For organoid-based regenerative therapy, AUF1-engineered organoids were administered to mice by rectal enema as described [51] with minor modifications. Organoids were harvested from Matrigel using gentle cell dissociation reagent, washed thoroughly with sterile PBS to remove residual extracellular matrix and culture medium, and resuspended in 100 µL of sterile PBS. At Day-7 of DSS administration, DSS-treated mice were lightly anesthetized, and the organoid suspension containing approximately 10^6^ organoid cells gently administered into the rectum using a flexible catheter. Control mice received an equivalent volume of PBS. After organoid transplantation, mice were subsequently monitored for disease progression and tissue regeneration.

### Statistical analysis

All data are presented as either mean ± standard error of the mean (SEM). Statistical analyses were performed using GraphPad Prism version 8.0.1 (GraphPad Software, San Diego, CA, USA). Unless otherwise indicated, data represent results from at least three independent experiments. For comparisons between two groups, a two-tailed unpaired Student’s *t*-test was used. Comparisons among more than two groups involving a single independent variable were performed using one-way analysis of variance (ANOVA) followed by Tukey’s post hoc multiple-comparisons test. For experiments involving two independent variables, two-way ANOVA followed by Šídák’s multiple comparisons test was performed. The number of biological replicates (*n*) and the specific statistical tests used for each experiment are indicated in the respective figure legends. A *P* value of <0.05 was considered statistically significant.

## Acknowledgements

We acknowledge the support of the Director, Indian Council of Medical Research–National Institute for Research in Bacterial Infections (ICMR-NIRBI), Kolkata, for facilitating this study. We express our deepest gratitude to Prof. Syamal Roy (IACS, Kolkata) for his support, valuable discussions, and helpful suggestions for the manuscript. We are grateful to all members of the Division of Immunology, ICMR-NIRBI, for their support. We also acknowledge Narayan Chandra Mondal for assistance with animal maintenance.

We acknowledge, Dr. Surojit Sinha, IACS, Kolkata for providing Morpholino oligomers for efficient gene knockdown. Additionally, we thank Dr. Jyotirmayee Dash, Senior Professor, School of Chemical Sciences, IACS, Kolkata, for assistance with the ITC experiments. The AUF1-EGFP isoform-containing constructs were kindly provided by Dr. Myriam Goropse, NIH while the AUF1-His_6_ constructs were kindly provided by Prof Gary Brewer, Rutgers University. We further acknowledge Prof. Dr. Suvendra Nath Bhattacharyya, IACS, Kolkata, for kindly providing the pre-miR-122 construct and Dr. Neeru Saini, IGBI, for providing the miR-195 construct.

OD is supported as an ICMR Project Research Scientist, and SAC is a recipient of a fellowship from the Department of Biotechnology (DBT), Government of India.

## Notes

### Competing Interest Statement

The authors have declared no competing interest.

