## Supplementary Figures for "AUF1-Engineered Intestinal Organoids Enhance Epithelial Barrier Repair and Mucosal Regeneration in Experimental Colitis"

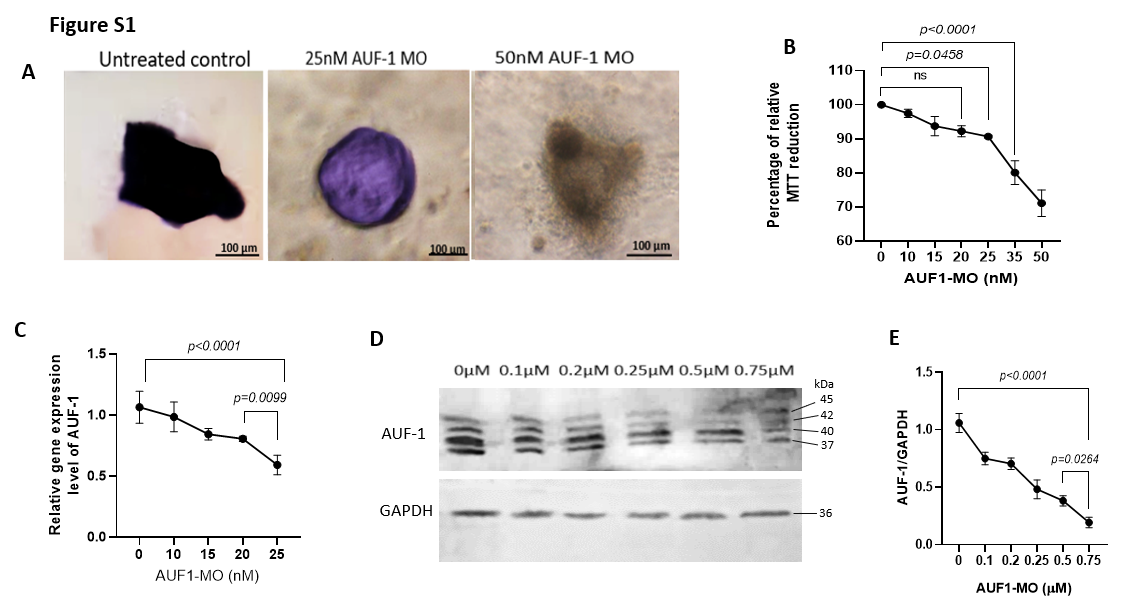

**Figure S1. Effect of AUF1-MO on organoid morphology, cell viability, and AUF1 expression. (A)**Representative images of organoids derived from control patients treated with or without 25 nM AUF-1 morpholino (AUF-1 MO), or 50 nM AUF-1 MO. Scale bars, 100 μm. **(B)** MTT assay showing the percentage of relative MTT reduction following treatment with increasing concentrations of AUF1-MO. **(C)** Relative AUF-1 gene expression following treatment with increasing concentrations of AUF1-MO. **(D)** Representative immunoblot showing AUF-1 protein levels following treatment with increasing concentrations of AUF1-MO, with GAPDH as the loading control. **(E)** Densitometric quantification of AUF-1 protein expression normalized to GAPDH. Statistical significance is indicated in the graphs.

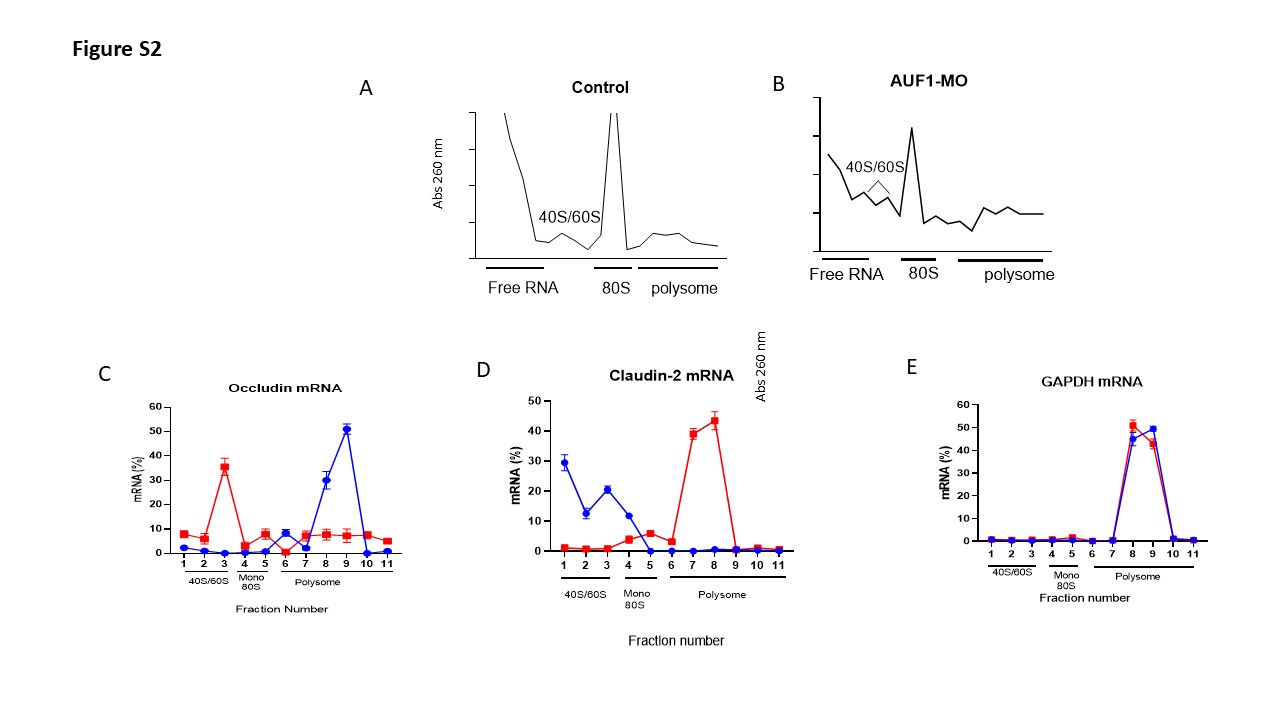

**Figure S2. Effect of AUF1-MO on polysome distribution and mRNA association.**

**(A–B)** Representative polysome profiles of control and AUF1-MO-treated samples, showing absorbance at 260 nm across free RNA, 40S/60S, 80S, and polysome fractions. **(C–E)** Distribution of **Occludin**, **Claudin-2**, and **GAPDH** mRNAs, respectively, across the indicated fractions. Fraction numbers and the corresponding 40S/60S, monosome/80S, and polysome regions are indicated below each graph. Error bars represent the variability shown for the individual measurements.

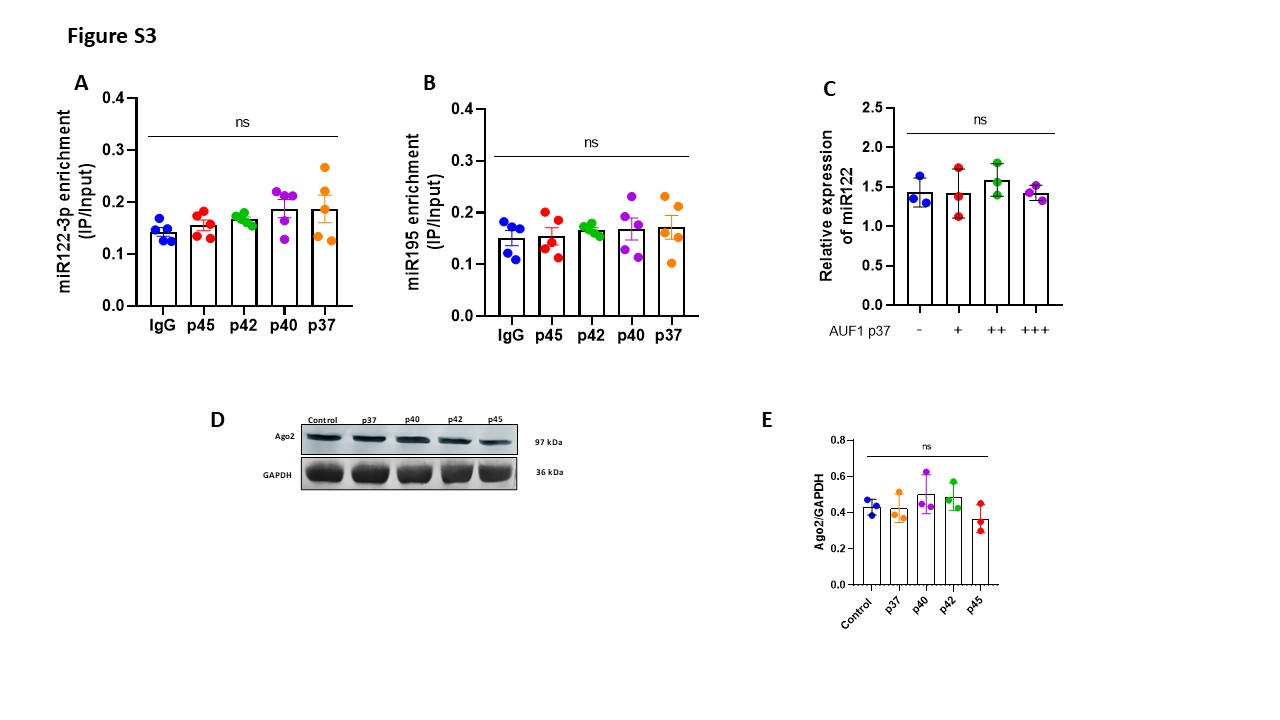

**Figure S3. Analysis of AUF1-associated miRNAs and Ago2 expression.**

**(A)**Enrichment of miR122-3p following immunoprecipitation with the indicated AUF1 isoforms compared with IgG control. **(B)** Enrichment of miR195 following immunoprecipitation with the indicated AUF1 isoforms compared with IgG control. **(C)** Relative expression of miR122 according to the indicated levels of AUF1 p37. **(D)** Representative immunoblot showing Ago2 protein levels in control and samples expressing the indicated AUF1 isoforms, with GAPDH as the loading control. Molecular-weight markers are indicated. **(E)** Densitometric quantification of Ago2 expression normalized to GAPDH. Statistical comparisons are indicated as shown; ns denotes not significant.

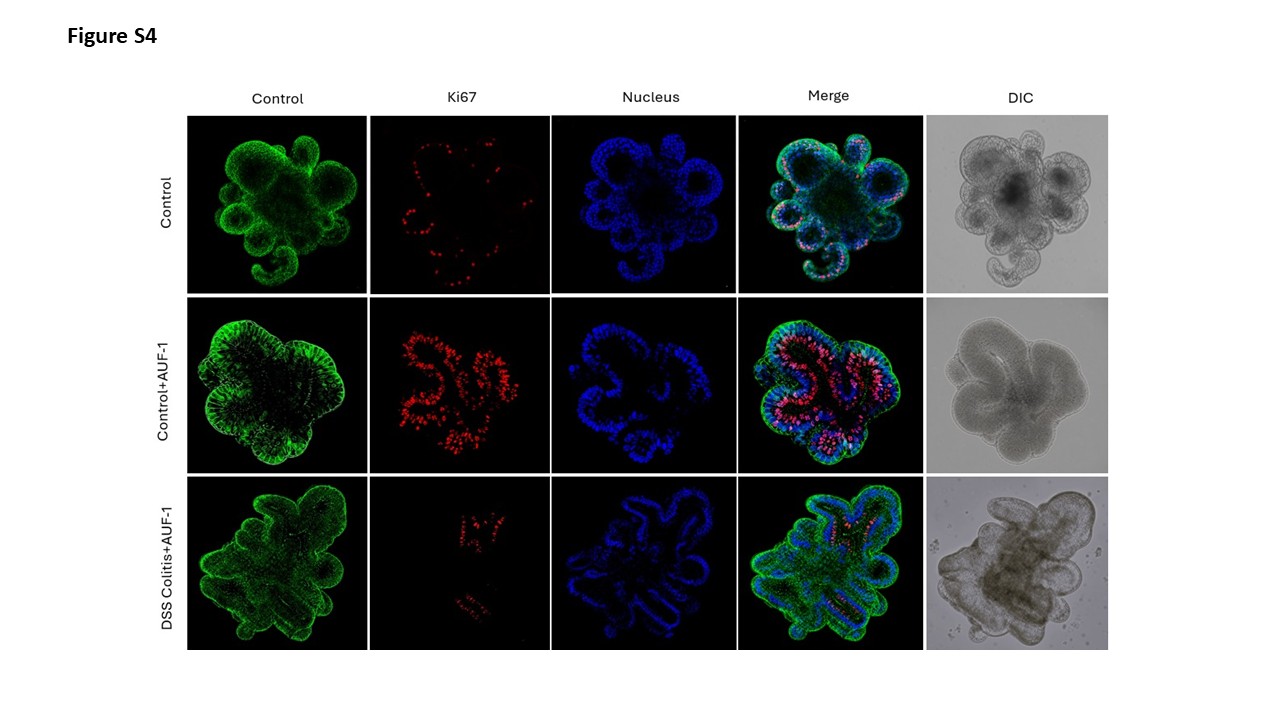

**Figure S4. Ki67 immunofluorescence analysis of organoids following AUF1 manipulation and DSS treatment.**

Representative images of organoids under the indicated experimental conditions: control, control transfected with EGFP- AUF-1p37+p40, and DSS colitis transfected with EGFP- AUF-1p37+p40. Images show the control/bright-field channel, Ki67 immunofluorescence, nuclear staining, merged fluorescence, and differential interference contrast (DIC) images. Ki67 staining is shown in red, nuclei in blue, and the control/organoid signal in green. The merged images demonstrate the spatial distribution of Ki67-positive cells within the organoids.

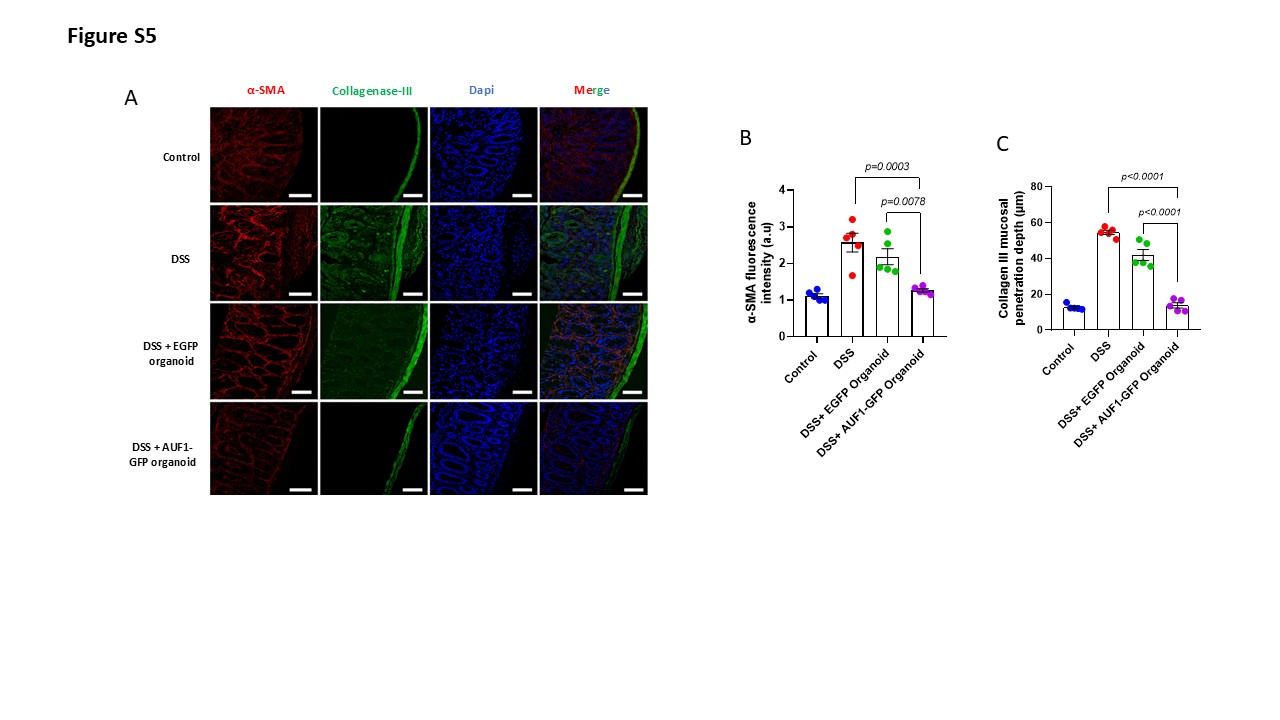

**Figure S5. Assessment of α-SMA expression and collagenase-III mucosal penetration in DSS-treated organoids.**

**(A)** Representative immunofluorescence images of control, DSS-treated, DSS + EGFP organoid, and DSS + EGFP-AUF1 organoid samples stained for α-SMA (red), collagenase-III (green), and nuclei with DAPI (blue); merged images are shown at right. Scale bars are shown in the images. **(B)** Quantification of α-SMA fluorescence intensity under the indicated experimental conditions. **(C)** Quantification of collagenase-III mucosal penetration depth (μm) in the indicated groups. Statistical significance is indicated above the corresponding comparisons.
